# Multi-modal choroid plexus pathology in aging and Alzheimer’s disease

**DOI:** 10.64898/2025.12.30.697051

**Authors:** Huixin Xu, Peter Lotfy, Benjamin Englert, Jane Oberhauser, Lillian I.J. Byer, Jasmin Wihlman, Kia Colangelo, Serhat V. Okar, Ashley Thommana, Henri Puttonen, Mikko I. Mäyränpää, Jarno Tuimala, Rute Pedrosa, Darshan Kumar, James Francis Haberberger, Micaiah Emmy Atkins, Shon Alimukhamedov, Aja Pragana, Jordan Benson, Mary Elizabeth Gabrielle, Anna Dong, Violeta Durán Laforet, Peter Bor-Chian Lin, C. Dirk Keene, Caitlin S. Latimer, Katherine E. Prater, David M. Holtzman, Alina Isakova, Tony Wyss-Coray, Dorothy P. Schafer, Daniel S. Reich, Terho Lehtimäki, Pekka J. Karhunen, Eloise H. Kok, Deidre Jansson, Andrew C. Yang, Liisa Myllykangas, Jose Ordovas-Montanes, Maria K. Lehtinen

## Abstract

Brain barriers, cerebrospinal fluid (CSF) dynamics, and peripheral factors are implicated as significant contributors to Alzheimer’s disease (AD). The choroid plexus (ChP) is a blood-brain interface that produces CSF and forms the blood-CSF barrier. However, how ChP pathology develops across the lifespan and contributes to AD has not been systematically characterized. Here, we report a multi-modal ChP atlas integrating single-nucleus transcriptomics from 49 individuals, AI-assisted quantitative histopathology across >500 postmortem samples age 16 to 105, spatial transcriptomics, and functional studies in 5xFAD mice. We identify fibrosis, calcification, and macrophage abnormalities as hallmarks of ChP aging, with AD pathology conferring additional effects, including expansion of a pro-inflammatory fibroblast-macrophage signaling niche. In 5xFAD mice, macrophage dysfunction is associated with impaired epithelial barrier maintenance and repair. Together, these data provide a foundational resource for understanding ChP dysfunction in aging and AD and propose the macrophage-fibroblast-epithelial barrier axis as a driver of ChP pathology.

## INTRODUCTION

The choroid plexus (ChP) is a secretory epithelial structure in the brain that produces cerebrospinal fluid (CSF) and forms the principal barrier between blood and CSF^1-3^. CSF has been extensively studied as a source of biomarkers for aging and Alzheimer’s disease (AD). In AD, changes in CSF amyloid-β and phosphorylated Tau levels predict disease up to 20 years before symptoms appear, and numerous additional biomarkers have since been identified^4-7^. Growing evidence also ties peripheral immune cells to brain aging and AD^8,9^, including accumulation of inflammatory immune cells in the CSF^10,11^. The ChP both monitors and actively modifies CSF composition through secretion and regulation of immune cell trafficking between the periphery and the central nervous system^12-16^. Therefore, AD-associated changes in CSF hint at possible ChP involvement beginning already from preclinical stages of AD. Clearance of brain waste products, including amyloid-β, through CSF circulation and drainage has emerged as a key contributor to brain homeostasis, and thus, ChP production of sufficient CSF would be cardinal to sustaining CSF flow and brain clearance^17^. However, despite the intimate connection between the ChP and CSF and emerging evidence linking ChP dysfunction to cognitive decline^18-25^, the ChP has received far less attention than the blood-brain barrier (BBB), which has been extensively characterized in neurodegeneration^26,27^. Together, these brain barriers protect the central nervous system from peripheral challenges; when compromised, they permit entry of blood-derived factors and immune cells that can facilitate neurodegeneration. While a “leaky” BBB is well established in aging and AD, the status and functional consequences of blood-CSF barrier integrity at the ChP remain incompletely defined.

The ChP is located within each brain ventricle and comprises convoluted layers of CSF-facing epithelial cells joined by tight junctions. The underlying stroma consists of fibroblasts, pericytes, and macrophages and is richly vascularized by fenestrated capillaries^1,28^. This architecture supports the dual function of the ChP as both a CSF secretory epithelium and a dynamic barrier between the CSF and blood that selectively gates the entry of blood-borne molecules into the CSF through regulation of tight junctions between epithelial cells^12-14^. This positions the ChP as a hub of body-brain communication whose roles in health and disease are only beginning to be understood. Pathological changes of the ChP are increasingly recognized in aging and neurological diseases^9,29-31^, from earlier studies in small cohorts of AD patients showing shortening of epithelial cells and thickening of basement membrane^32^, to inflammatory changes in amyotrophic lateral sclerosis^33^, fibrosis in multiple sclerosis^34^, and formation of Biondi bodies in aging^35,36^. Furthermore, MRI studies consistently report ChP enlargement with age and have identified associations with a number of neurodevelopmental and neurodegenerative disorders^30^, including AD. Multiple independent studies have identified an association between ChP enlargement and AD-relevant traits: total ChP volume correlates with cognitive decline, hippocampal shrinkage, and dysfunction of CSF dynamics, while the rate of enlargement is exacerbated by risk factors including female sex and APOE ε4 genotype^19,37-40^. These structural changes are hypothesized to impair ChP function through several proposed mechanisms, including barrier leakage permitting entry of blood-derived factors and immune cells into the CSF, reduced CSF secretion and turnover, and consequently, insufficient clearance of waste and neurotoxic proteins such as amyloid-β.

Recent multi-omics studies of human and mouse ChP in AD point towards transcriptional and proteomic changes with implications for ChP epithelial dysfunction and inflammation^41-43^. However, the cellular and molecular changes underlying ChP dysfunction across the lifespan have not been systematically defined, precluding identification of specific pathological mechanisms and therapeutic opportunities. Which ChP cell types are affected, what molecular programs drive pathological changes, and whether aging and AD exert separable or synergistic effects remain open questions. Addressing these questions is essential to understanding whether ChP dysfunction represents a cause or consequence of AD and whether it constitutes a tractable therapeutic target.

Here, we generated a multi-modal atlas of ChP pathology in aging and AD, integrating studies of human postmortem specimens and mouse models to distinguish age-associated changes from disease-specific effects. We combined single-nucleus RNA-sequencing (snRNA-seq, 49 patients from two cohorts, age range 70-101 years old), 10X Xenium spatial transcriptomics (1 male patient, age 48 years old), Artificial intelligence (AI)-assisted quantitative histopathology (529 patients, age range 16-105 years old), and patient magnetic resonance imaging (MRI) scans (20 patients, age range 31-62 years old). This approach identified extensive fibrosis, calcification, and myeloid cell abnormalities as hallmarks of ChP aging, with sex-dependent effects of AD pathology. We further used a well-established mouse model of amyloid deposition in AD (5xFAD^44^) to dissect ChP changes at early-to-mid stages of AD pathology, independent of aging. Longitudinal *in vivo* two-photon imaging revealed dysregulated macrophage number, morphology, behavior, and recruitment linked to deficits in epithelial barrier maintenance and repair. Together, these data provide a foundational resource for understanding ChP dysfunction and nominate the macrophage-fibroblast-epithelial barrier axis as a candidate driver of ChP pathology in aging and neurodegeneration.

## RESULTS

### A cellular atlas of the choroid plexus in AD reveals disease-associated inflammatory states

To systematically characterize ChP cellular changes in aging and AD, we generated a snRNA-seq atlas of 197,108 nuclei from postmortem lateral ventricle ChP collected from 49 individuals from two independent cohorts: the Religious Orders Study/Memory and Aging Project (ROSMAP; n=31) and the University of Washington Alzheimer’s Disease Research Center (ADRC; n=18; **Figure 1A, Supplemental Data 1**). These cohorts included individuals with pathology-confirmed AD (n=21), mild cognitive impairment due to AD (MCI; n=10), and cognitively normal controls without significant neuropathology (n=18) and were balanced for age, sex, APOE genotype, and post-mortem interval (**Figure S1A**). Clinical and neuropathological metadata, including Braak staging and CERAD scores, were not significantly correlated with demographic metrics, including sex, age, or post-mortem interval (**Figure S1B**). Disease and age:disease were shown to be the primary drivers of variance in the dataset, followed by sex and age:sex (**Figure S1D**).

**Figure 1.**
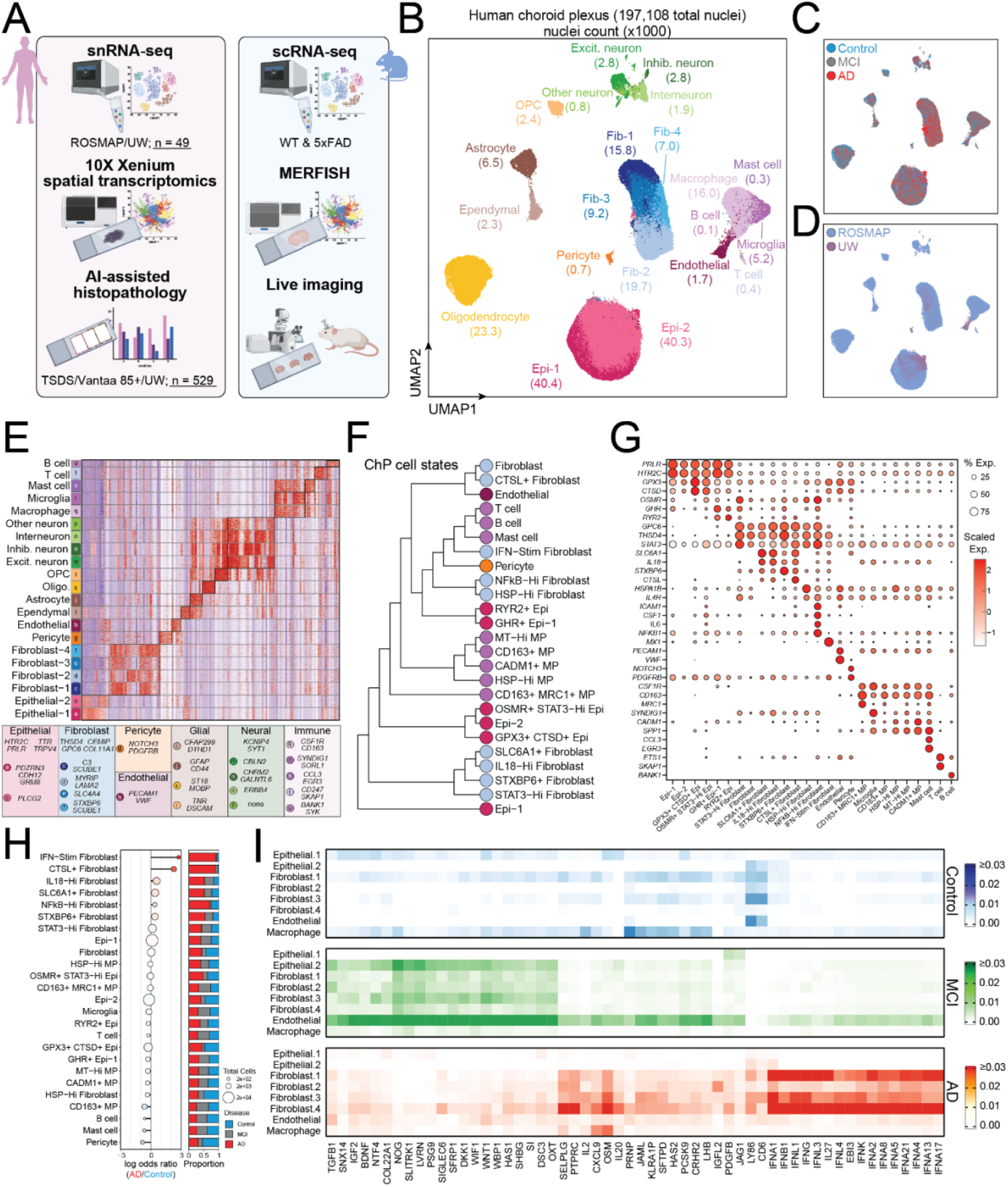
A cellular atlas of the choroid plexus in AD reveals widespread inflammatory states. (A) Schematic overview of the multi-modal study design. Human ChP was profiled using snRNA-seq, 10X Xenium spatial transcriptomics, and AI-assisted histopathology (TSDS/Vanta85+/UW, n=529 donors). Mouse ChP was profiled using scRNA-seq, MERFISH spatial transcriptomics, and *in vivo* two-photon imaging in WT and 5xFAD mice. (B) Uniform Manifold Approximation and Projection (UMAP) embeddings of 197,108 nuclei (49 donors) isolated from human ChP colored by cell type annotations. Numbers indicate cell counts (x1000) per cell type. (C) UMAP colored by disease classification (Control: blue, mild cognitive impairment (MCI): grey, AD: red). (D) UMAP colored by study of origin (ROSMAP: blue, n=168,171, 31 donors; UW: purple, n=28,937, 18 donors). (E) Heatmap of marker gene expression across cell subtypes. Color scale represents gene expression as a z-score (navy: low, white: intermediate, red: high). Bottom panel: key marker genes organized by cell class. (F) Hierarchical clustering dendrogram of ChP cell states based on transcriptomic similarity. Clustering was performed using Ward’s method on Euclidean distances of scaled average expression of variable genes. MP: macrophage; Epi: epithelial cells. (G) Dot plot of marker gene expression across cell states. (H) Compositional analysis of cell state abundance in AD versus Control. Left: Proportional regression analysis showing log odds ratio with Benjamini-Hochberg FDR values of enrichment of a cell state in AD (red) compared to Control (blue). Right: stacked bar plots showing disease group proportions for each cell state. (I) Heatmaps of predicted MultiNicheNet ligand activities across disease conditions. Only ligands predicted to activate target genes are shown.

Following quality control and batch correction with Harmony^45^ (**Figure S1C-D**), unsupervised clustering of the integrated dataset identified 14 major cell type clusters. Eight represented canonical ChP-resident and hematopoietic cell types, including endothelial cells, epithelial cells, fibroblasts, pericytes, macrophages, mast cells, T cells, and B cells. The other six, including neurons, astrocytes, ependymal cells, oligodendrocytes, oligodendrocyte precursor cells (OPCs), and microglia, were likely captured during tissue dissection due to the anatomical proximity of the ChP to cortical tissue (**Figure 1B-C**). Gene set enrichment analysis confirmed expected functional profiles: epithelial cells were enriched for ion channel transport, fibroblasts for extracellular matrix (ECM) remodeling, and pericytes for muscle contraction (**Figure S1E**).

To define the cellular heterogeneity of the human ChP at higher resolution, we further subclustered these ChP populations hierarchically at two levels (**Figure 1E-G**). First, we subclustered into 21 clusters across all subtypes, revealing further diversity in epithelial cells (2 clusters), fibroblasts (4 clusters), and neurons (4 clusters). Epithelial-1 cells were defined by elevated expression of *PDZRN3*, *CDH12*, and *GRM8*, while Epithelial-2 cells were defined by *PLCG2*. Fibroblast-1 cells expressed Complement C3 (*C3*) and *SCUBE1*, Fibroblast-2 expressed contractility genes *MYRIP* and *LAMA2*, Fibroblast-3 expressed the bicarbonate transporter *SLC4A4*, and Fibroblast-4 cells were defined by their expression of *STXBP6* (**Figure 1B-E**). Following exclusion of glial and neural cells, we further subclustered to identify 26 clusters representing transcriptionally distinct cell states (**Figure 1F**). This revealed several clusters of epithelial cells and fibroblasts expressing high levels of inflammatory response genes, including epithelial cells expressing high levels of Oncostatin M receptor (*OSMR*) and its downstream effector *STAT3*, which promotes inflammatory gene expression in response to IL-6 family cytokines. Among fibroblasts, we observed STAT3-high Fibroblasts co-expressing the STAT3 target gene *IL4R*^46^, interferon-stimulated genes (*MX1*, *ISG15*), and NF-κB-activated fibroblasts expressing *CSF1*, *IL6*, *ICAM1*, and *NFKB1*. The presence of these inflammatory states across multiple ChP cell types suggests coordinated inflammatory signaling within the tissue microenvironment.

To determine if these states were enriched in AD, we conducted differential abundance analysis of cell states between AD and control samples. This revealed high enrichment of multiple inflammatory fibroblast states in AD samples, including Cathepsin L (*CTSL*)-positive and IFN-stimulated fibroblasts, though these did not reach statistical significance after multiple hypothesis correction (**Figure 1H**). These trends were instead driven by elevated frequencies in a subset of AD patients rather than a consistent shift across the cohorts, reflecting substantial heterogeneity in fibroblast inflammatory responses among AD individuals. This patient-level variability is consistent with the known clinical and pathological heterogeneity of AD^21^ and provides further evidence that fibroblast inflammatory activation may characterize a subset of AD patients rather than a universal disease feature.

To infer the upstream signaling pathways driving these transcriptional states, and how they may differ between disease classifications, we conducted ligand activity inference using MultiNicheNet^47^ (**Figure 1I**). We observed significant disease-enriched type I and type II interferon (IFN) signaling in fibroblasts and disease-enriched NF-κB signaling in epithelial cells. Furthermore, we found predicted STAT3-related cytokine activity, including Oncostatin M (*OSM*) and IL-20 in the stroma. Together, these findings establish a comprehensive cellular atlas of the human ChP and reveal inflammatory programs that implicate interferon, NF-κB, and STAT3 signaling in ChP dysfunction during AD progression.

### AI-assisted histopathology identifies fibrosis and calcification as age-driven ChP pathologies

Given the inflammatory signatures observed in fibroblasts and epithelial cells, we next investigated cellular pathologies using >500 human pathology specimens. We used AI-assisted quantitative histopathology in three independent cohorts of patient specimens to both determine AD pathology-driven versus age-driven changes, focusing on reported hallmarks including fibrosis, psammoma bodies (calcification), and inflammation^30,32^. The first cohort, the Vantaa 85+ study, is a population-based cohort consisting of all individuals aged >85 years living in Vantaa, Finland on April 1, 1991^48-50^ (n=282; **Supplemental Data 2**). The second cohort, the Tampere Sudden Death Study (TSDS), is a forensic autopsy series comprising white Caucasians^51-53^ (n=181; age range 16-90; **Supplemental Data 2**). The third cohort was collected from University of Washington (UW) through Alzheimer’s disease research center (ADRC) (n=66; **Supplemental Data 2**).

Fibrosis and psammoma bodies (calcification) were found to correlate positively with aging in the ChP (**Figure 2A-B**). Masson’s trichrome staining was applied to ChP samples to detect connective tissue (collagen) and identify fibrotic ChP villi. We trained an AI model (Aiforia Technologies) to detect ChP tissue area, normal villi, fibrotic villi with increased deposition of connective tissue (**Figure 2A**), and psammoma bodies shown as concentric rings of calcium deposits that accumulated especially within degenerative areas (**Figure 2C**). Fibrosis was calculated by dividing the number of fibrotic villi by the number of normal villi. The number of psammoma bodies was normalized by the tissue area. Both ChP fibrosis and calcification correlated with age. We analyzed fibrosis and psammoma bodies in association with age and various AD pathologies (**Supplemental Data 2**). In the TSDS cohort, fibrosis (p≤0.001) and psammoma bodies (p≤0.023) both increased with age. In the Vantaa 85+ cohort, where all subjects were over 85 years old, fibrosis showed a borderline significant association with Thal phases as AD-related pathology (p≤0.054) and psammoma bodies trended (p≤0.066) with age in males. In the UW cohort, age- and sex-adjusted analyses showed that psammoma bodies were associated with CERAD (p≤0.022) and NIA-ABC scores (p≤0.046). Collectively, our data from multiple independent large cohorts of human samples demonstrated strong age-driven ChP fibrosis and calcification. AD pathology showed moderate association after age-adjusted analyses with some sex-dependence.

**Figure 2.**
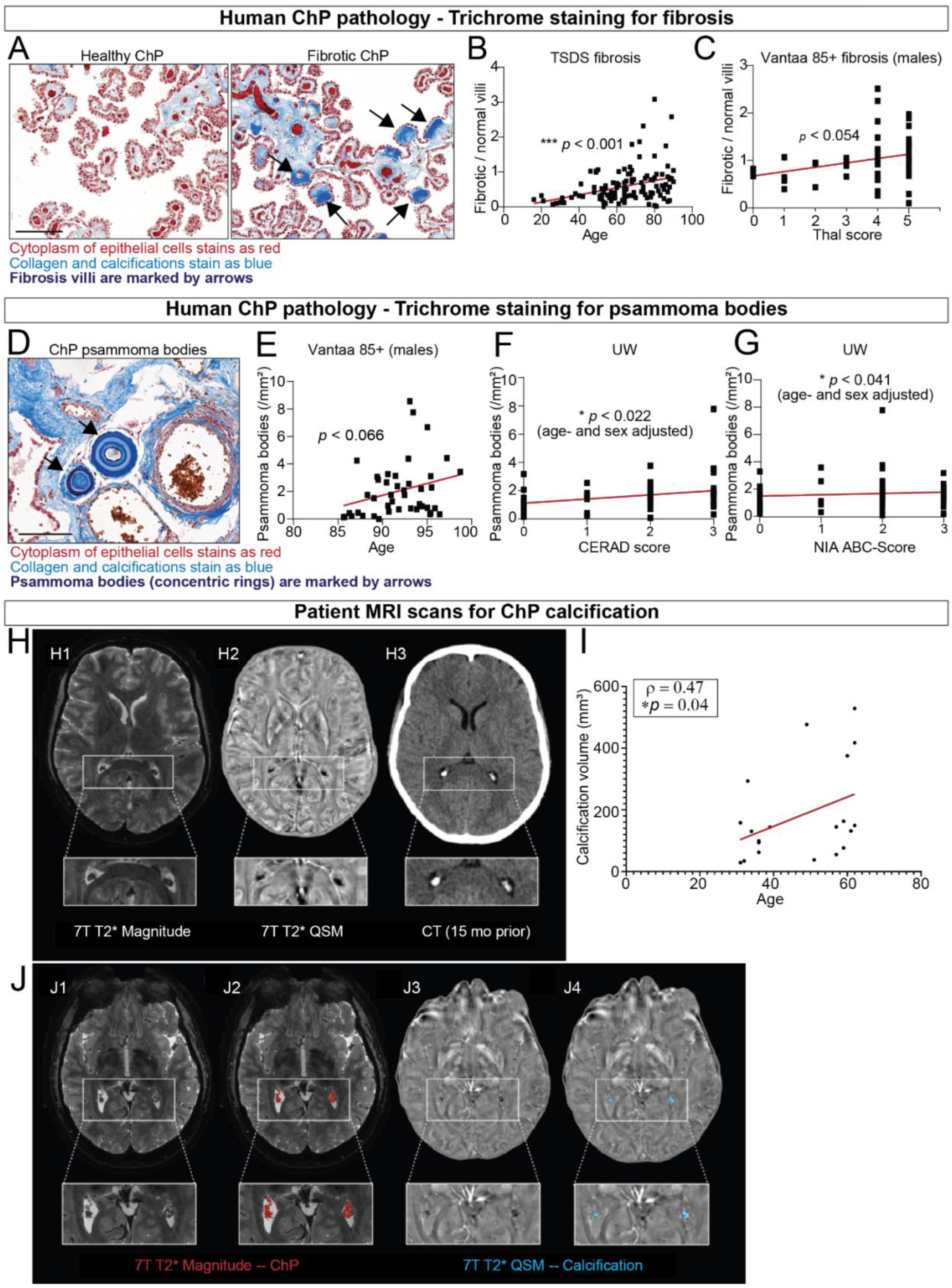
AI-assisted histopathology identifies fibrosis and calcification as age-driven ChP pathologies in human. (A) Representative images showing trichrome stained human ChP sections from aged subjects. Condensed dark blue staining indicates fibrosis (marked by arrows). Scale = 300 µm. (B) Quantification by Aiforia showing the ratio of fibrotic versus healthy villi is associated with age in TSDS cohort (trichrome staining, simple linear regression model; p < 0.001). (C) Ratio of fibrotic per normal villi (quantified by Aiforia) in relation to Thal β-amyloid phases in males of the Vantaa 85+ cohort (trichrome staining, simple linear regression model; p < 0.054) (D) Representative images showing trichrome stained human ChP sections from aged subjects. Blue/reddish concentric lamellar/roundish structures indicate calcification (marked by arrows), in perivascular location. Scale = 300 µm. (E) Psammoma bodies (/mm^2^) quantified by Aiforia on trichrome stainings in relation to age in males (Vantaa 85+, simple linear regression, p < 0.066). (**F, G**) Aiforia quantified Psammoma bodies (/mm^2^) in the UW cohort are associated with CERAD (F; p < 0.022) and NIA-ABC (G; p < 0.041) scores (trichrome staining, linear regression adjusted for age and sex). (H) Representative images of patient MRI and CT scans. (D1) Choroid plexus from a 34-year-old woman with MS visualized with 7T T2* GRE. (D2) 7T T2* QSM demonstrated hypointense regions that are calcified, aligning with CT findings showing hyperdense calcified regions (D3). (I) Quantification of patient MRI scans showing calcification of the ChP is associated with age. (J) Representative images showing the segmentation of the ChP and calcification. (J1) Choroid plexus and the corresponding segmentation (J2) on 7T T2* GRE from a 31-year-old man with MS. (J3) Calcified hypointense regions on 7T T2* QSM were identified, and corresponding segmentations (J4) were created.

To validate these findings using orthogonal imaging modalities, we applied Micro-CT imaging of ChP specimens from Vantaa 85+ cohort, confirming age-associated ChP calcification (*p* < 0.045) (**Supplemental Data 2**). Analysis of brain MRI scans from an independent cohort (n=20 multiple sclerosis patients receiving MRIs) revealed that calcification of the lateral ventricle ChP correlated with age but not disease parameters and was notably more extensive in the ChP than in other brain regions. These data suggest that the ChP is particularly vulnerable to age-related calcification (**Figure 2D-F**).

To determine the cellular basis of ChP dysfunction and microanatomical heterogeneity in AD, we performed Xenium spatial transcriptomics on the ChP of a 48-year-old, cognitively normal male donor (**Figure S2A-F**), enabling identification of direct cell contacts in the ChP. The spatial data recapitulated all ChP cell types identified in our snRNA-seq dataset and revealed distinct microanatomical localization and spatial niches of different cell subtypes in the ChP stroma, including four subtypes of fibroblasts (**Figure 3A-B**). We found that fibroblasts and macrophages differentially but also preferentially co-localize across ChP spatial niches (**Figure 3B-E, Figure S2G)**. Using ligand-receptor interaction analysis in our snRNA-seq dataset, we found that several putative pro-fibrotic and pro-inflammatory fibroblast-macrophage interactions were enriched in AD compared to control, including transforming growth factor β (TGF-β) signaling from macrophages to fibroblasts and collagen-associated factors from fibroblasts to macrophages (**Figure 3F**). Previous work from our group and others has shown that local sources of M-CSF (encoded by *CSF1*) are essential for the maintenance of tissue macrophages^14,54^. We found here that inflammatory fibroblasts are the primary source of CSF1 in the human ChP (**Figure 3G**), suggesting that inflammatory fibroblasts in the ChP promote the retention and recruitment of pro-fibrotic macrophages in AD. Together, these data illustrate that ChP macrophages display a regional, niche-dependent predilection for close physical interaction with ChP fibroblast subtypes and other macrophages or immune cells, suggesting the possibility of a proximity-driven role in later age or disease-related inflammatory processes.

**Figure 3.**
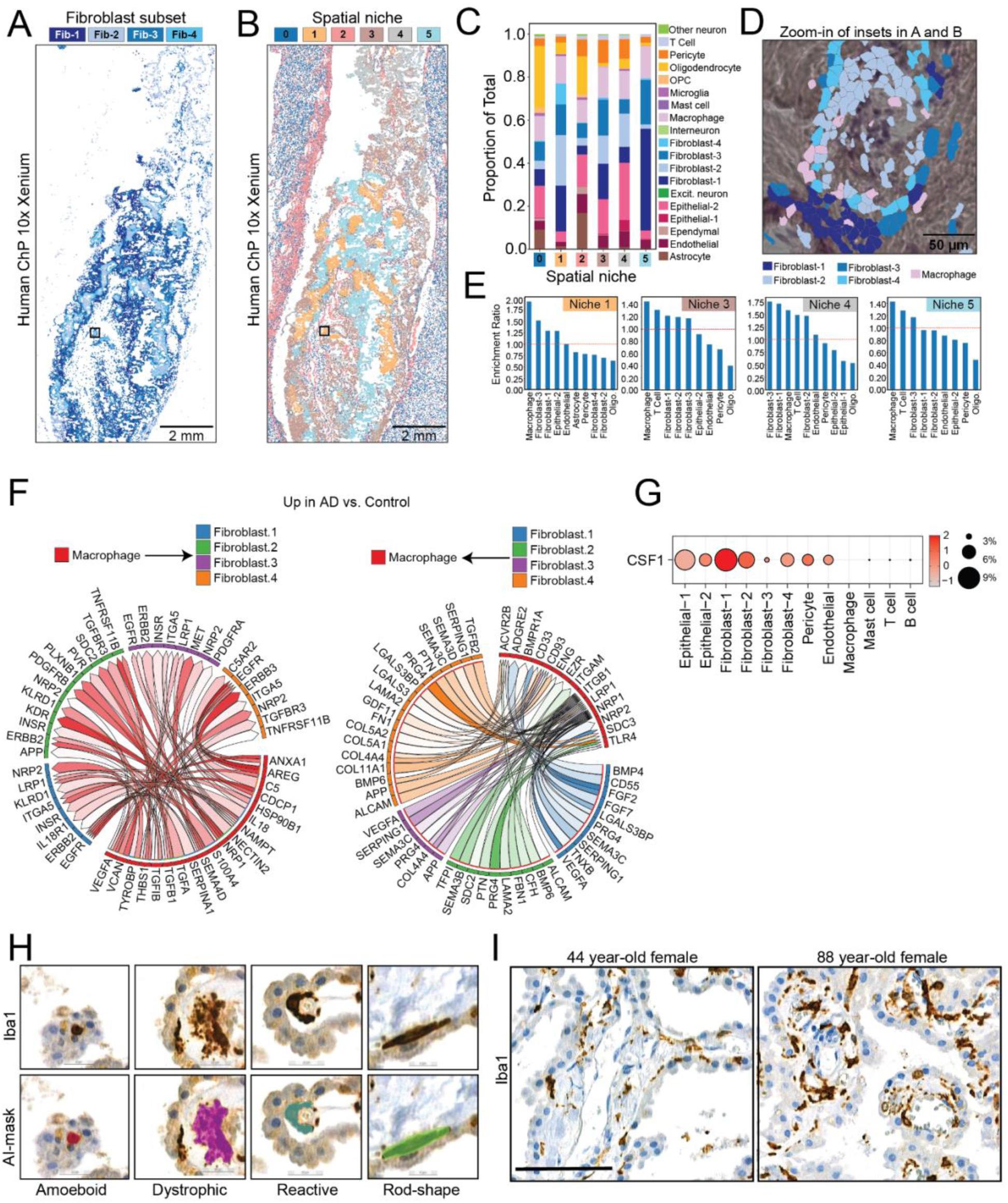
Spatial and single-nucleus transcriptomics identifies fibroblast-macrophage niches in human ChP. (**A**) Xenium cell type distributions reveal the differential localization of functionally varied choroid plexus fibroblast subtypes identified by snRNA-seq. (**B-C**) Graph embedding defines choroid plexus niches defined by distinct spatial distributions, anatomical characteristics, and cell type proportions. Scale bar: 2 mm. Black box indicates region shown in (D). (D) Spatial co-localization of choroid plexus macrophages and various fibroblast subtypes within spatial niche 1. Scale bar: 50 µm. (E) Choroid plexus macrophages display niche and region-specific interactions with fibroblast subtypes, epithelial cells, and other immune cells based on the relative frequency of cell-cell proximity. (F) Chord diagram depicting top 50 ligand-receptor interactions upregulated in AD versus Control (log2FC > 0.50 and p<0.05) between macrophages and fibroblast subtypes. (G) Dot plot of *CSF1* expression across ChP cell types. (H) Representative images showing Iba1+ macrophages of various morphology and their masks generated by Aiforia for quantification. (I) Representative images showing increased Iba1+ macrophage coverage in the ChP in aged humans. Scale = 100.

Because of this observation and the well-established macrophage-fibroblast circuit^54^, we sought out to explore whether a pro-inflammatory fibroblast-macrophage interaction could promote inflammation and ChP dysfunction in aging and AD. To quantitatively analyze macrophage number and inflammation-associated morphological changes, we trained the Aiforia classifier to categorize Iba1+ macrophages by morphology (**Figure 3H**). In Vantaa 85+ cohort, total macrophage coverage (p≤0.003, **Figure 3I**) and fraction (p≤0.009) of reactive macrophages associated with age. In men, a positive association between Braak stages (controlled for age) and the fraction of amoeboid macrophages (p≤0.032), a morphology indicative of a more reactive, pro-inflammatory phenotype, was found. In the UW cohort, an age-association could be seen for the coverage (p≤0.023) and fraction (p≤0.021) of rod-shaped macrophages. All other analyses between macrophage morphology and AD pathology or age can be found in **Supplemental Data 2**. Collectively, our analyses revealed strong age-driven changes in macrophage morphology consistent with increased inflammation. AD pathology showed moderate additional association with some sex-dependence, similar to our observation in fibrosis.

While human histopathology revealed robust age- and sex-associated changes in the ChP, the correlation between age and AD pathology in humans limits causal inference. To disentangle these factors and analyze ChP changes in response to a broad range of AD pathologies from mild to severe, we turned to mouse models.

### Mouse models of AD reveal abnormalities in macrophage behaviors and barrier structure

To isolate AD-associated pathology from normal aging and gain more mechanistic insights, we analyzed the ChP of 5xFAD mice, a mouse model of Aβ amyloidosis, with males and females equally included and analyzed separately, across disease progression (2-9 months of age). Because 5xFAD mice have accelerated AD pathology, we did not detect any fibrosis by 9 months of age, which supports the human pathology findings that aging is the main driving force for ChP fibrosis. Instead, we found inflammatory changes occur at early-to-severe-stages of AD-like pathology, some of which appeared prior to extensive plaque deposition. We first characterized ChP macrophages and their responses to early disease pathology. ChP macrophages occupy two distinct anatomical compartments (**Figure S3A**): the apical, CSF-contacting surface of the ChP (epiplexus) and the underlying stromal layers located between epithelial cells and blood vessels (stromal). We previously demonstrated that epiplexus macrophages respond to CSF-derived stimuli, whereas stromal macrophages surveil blood-derived factors^55^. AD pathology alters CSF composition in both human and mouse models, including elevated neurofilament light chain (NfL) and altered Aβ42/40 ratios^4,56,57^. Given their strategic positioning, ChP epiplexus macrophages are well-positioned to sense and respond to these pathological changes. As early as 2 months of age, 5xFAD mice exhibited a striking increase in epiplexus macrophage density compared to wild-type (WT) controls (**Figure 4A-B**), and this elevation persisted at 6 months (**Figure 4C**). At 2 months, 5xFAD mice also showed increased total macrophage coverage (measured by total CX3CR1+ area in wholemount explants) (**Figure 4D**) and longer average process length (**Figure 4E**). By 6 months, total coverage and average process length no longer differed significantly between genotypes; however, 5xFAD mice displayed a subpopulation of epiplexus macrophages with markedly long processes (>75 µm compared to <25 µm typical of WT) as well as a subpopulation of macrophages lacking processes entirely (**Figure 4F**), suggesting increased morphological heterogeneity with disease progression.

**Figure 4.**
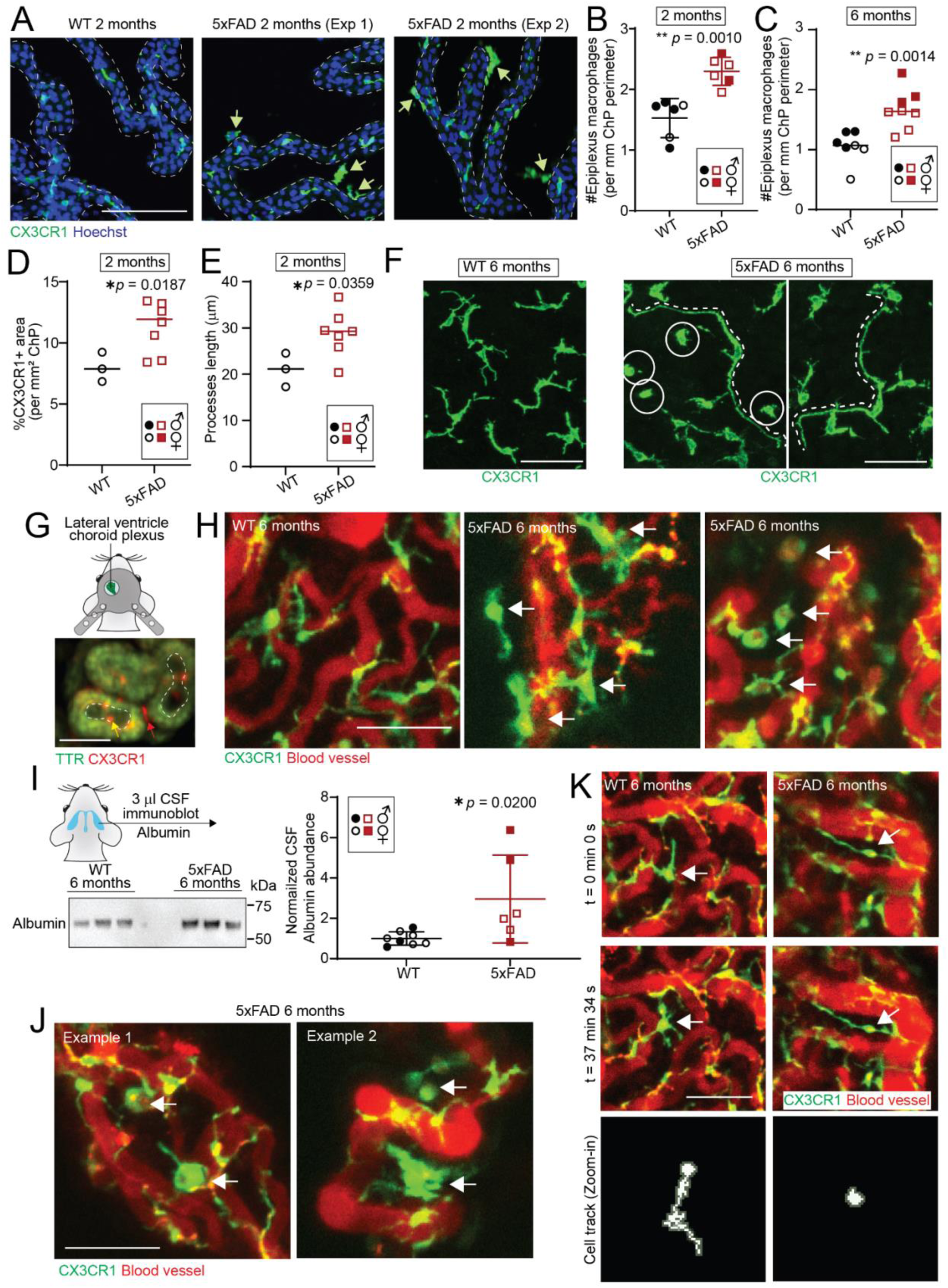
Mouse models of AD reveal abnormalities in macrophage behaviors. (A) Representative images showing increased number of epiplexus macrophages in the ChP of 5xFAD mice as early as 2 months old. Scale = 100 µm. (**B-C**) Quantification showing the number of epiplexus macrophages per mm of ChP perimeter is higher in 5xFAD mice than WT controls at both 2 months (B, ** p = 0.0010) and 6 months old (C, ** p = 0.0014). Welch’s t-test. (D) Quantification showing the total area of CX3CR1+ macrophages per mm^2^ of ChP increases in 5xFAD mice at 2 months old. * p = 0.0187. Welch’s t-test. (E) Quantification showing the average process length of CX3CR1+ ChP macrophages increases in 5xFAD mice at 2 months old. * p = 0.0359. Welch’s t-test. (F) Representative images showing ChP macrophages with extremely long processes and no processes present in 5xFAD mice at 6 months. Scale = 50 µm. (G) Schematics and a representative image from a TTR^mNeon^/CX3CR1^TdTomato^ mouse showing *in vivo* visualization of the ChP through a cranial window and separation of epiplexus versus stromal macrophages. Epiplexus macrophages (red arrow) are seen on top of epithelial cells (Green, TTRmNeon+) with a clear distance to blood vessels (dotted outline), while stromal macrophages (yellow arrow) are immediately adjacent to blood vessels. Scale = 50 µm. (H) Representative snapshots of in vivo recordings showing WT ChP (left) versus two example FOVs of 5xFAD ChP that contain increased number of epiplexus macrophages (middle, white arrows) and ameboid epiplexus macrophages with red blood-borne dye (right, white arrows). Scale = 100 µm. (I) Representative blot and quantification showing increased CSF albumin levels in 6 months old 5xFAD mice. * *p* = 0.0200. Mann-Whitney. (J) Representative snapshots of *in vivo* recordings showing two examples of macrophages with large vacuoles (white arrows) assembling phagocytic bubbles. Scale = 100 µm. (K) Representative snapshots of *in vivo* recordings at the beginning and the end showing lack of movement in the 5xFAD macrophage with extremely long processes. Scale = 100 µm. The tracks of the cell body of the chosen macrophages (white arrows) through the entire recordings are displayed at the bottom (with zoom-in for clarity). All quantitative data are presented as mean ± SD. Female mice are marked by unfilled legends.

*In vivo* two-photon imaging revealed distinct macrophage behaviors in 5xFAD mice compared to WT controls. We crossed 5xFAD heterozygous males with *Cx3cr1^GFP^* reporter females to generate 5xFAD;*Cx3cr1^GFP^* and WT;*Cx3cr1^GFP^*littermates for imaging at 6 months of age. Following recovery from cranial window implantation, we administered 70 kDa Texas Red-conjugated dextran intraperitoneally 15 minutes prior to imaging to label blood vessels. Epiplexus and stromal macrophages were distinguished based on their position relative to blood vessels (**Figure 4G**). As illustrated in mice with epithelial cells labeled by *Ttr^mNeon^*, epiplexus macrophages are visible on top of epithelial cells at a clear distance from blood vessels ( **Figure 4G**, outlined), while stromal macrophages lie beneath the epithelium near the blood vessels^14^. Each mouse was recorded across two fields of view for approximately one hour per day over two non-consecutive days. Live imaging revealed several abnormal phenotypes from CX3CR1+ ChP macrophages in 5xFAD mice. First, 5xFAD ChP contained focal areas with increased numbers of enlarged epiplexus macrophages (**Figure 4H**, middle; **Supplemental Video V1**) compared to WT controls (**Figure 4H**, left; **Supplemental Video V1**). Second, we observed ameboid epiplexus macrophages containing intracellular dextran reflecting uptake of blood-borne dextran (**Figure 4H**, right; **Supplemental Video V1**). This intracellular accumulation of dextran may reflect focal epithelial barrier leakage, which was further supported by increased levels of albumin, a blood-derived protein, in the CSF of 5xFAD mice (**Figure 4I**). Consistent with enhanced phagocytic behavior, we also captured epiplexus macrophages containing large cytoplasmic vacuoles resembling those observed in LPS-stimulated cells (**Figure 4J, Figure S3B**). Third, at baseline, epiplexus macrophages are highly mobile with continuously moving cell bodies^55^; however, 5xFAD mice harbored epiplexus macrophages with extremely long processes that appeared anchored to the epithelium (**Figure 4K, Supplemental Video V2**). These cells exhibited markedly reduced motility, with their cell bodies showing minimal displacement over 30 minutes compared to the nearly continuous and dynamic repositioning observed in WT controls (**Figure 4K, Supplemental Video V2**). These altered morphologies and impaired motility suggest dysregulated macrophage function in 5xFAD mice, potentially associated with compromised barrier integrity.

In further support of ChP barrier damage, we observed macrophages clustering near blood vessels in 5xFAD mice. Both whole-mount immunofluorescence and *in vivo* imaging revealed approximately 1-3 distinct macrophage clusters per lateral ventricle ChP in 5xFAD mice, extending from the apical CSF-facing surface through the stroma to the adjacent blood vessels (**Figure 5A-C, Supplemental Video V3**). Strikingly, these clusters were rarely observed in age-matched WT controls. A subset of cells within these clusters expressed high levels of CD45 (**Figure 5B**), consistent with recently infiltrated peripheral myeloid cells. The perivascular aggregation of myeloid cells resembles patterns we previously observed in the ChP during laser-induced micro-hemorrhage repair^55^, suggesting these clusters may represent sites of vascular damage and active repair.

**Figure 5.**
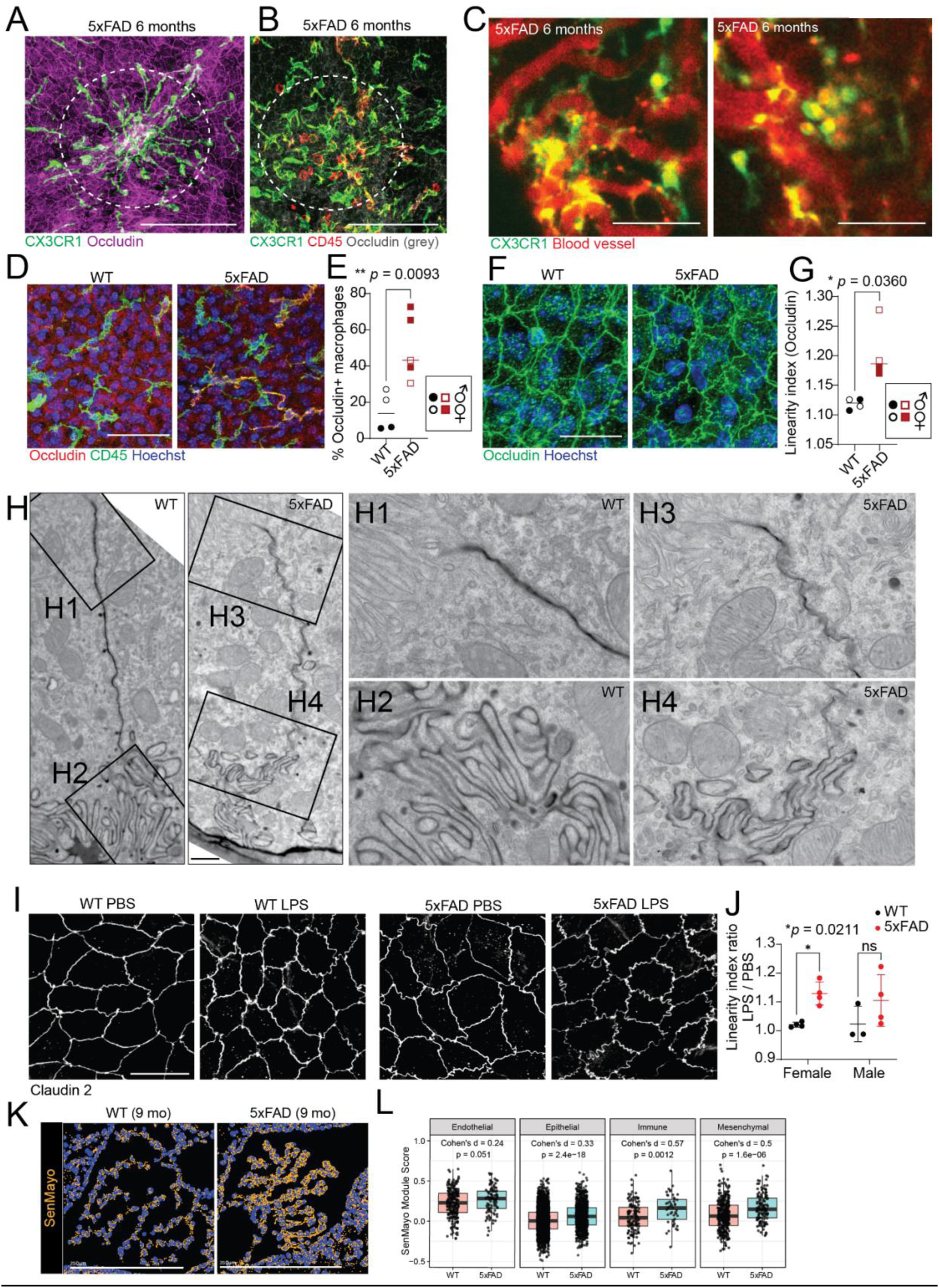
Macrophages are involved in epithelial barrier dysregulation in mouse models of AD. (**A-B**) Representative images showing clusters of CX3CR1+ macrophages and CD45-high peripheral immune cells at the ChP of 6 months old 5xFAD mice. Scale = 100 µm. (**C**) Representative snapshots of in vivo recordings showing two examples of macrophages clustering near blood vessels in the ChP of 6 months old 5xFAD mice. Scale = 100 µm. (**D-E**) Representative images and quantification showing increased portion of ChP macrophages stain positive for tight junction protein Occludin in 5xFAD mice at 6 months. Scale = 100 µm. ** *p* = 0.0093. Welch’s t-test. (**F-G**) Representative images and quantification showing increased linearity index of epithelial cell-cell borders marked by tight junction protein Occludin, indicating barrier damage. Scale = 50 µm. * *p* = 0.0360. Welch’s t-test. (**H**) Representative images of electron microscopy of ChP epithelial cells showing “wavy” cell-cell borders in 6 months old 5xFAD mice. Cell-cell borders are labeled by retro-orbital infusion of HRP. H1 and H3 show the zoom-in views at the apical surface. H2 and H4 show the zoom-in views of the basal labyrinth. Scale = 1 µm. (**I-J**) Representative images and quantification showing delayed epithelial barrier recovery in female 5xFAD mice at 6 months old, shown by sustained high linearity index of cell-cell borders labeled by tight junction protein Claudin 2. Scale = 20 µm. * *p* = 0.0211. Welch’s t-test. (**K-L**) Spatial transcriptomic analysis of 9 months 5xFAD mice ChP by MERFISH reveals that 5xFAD mice ChP had increased senescence score (SenMayo) in major cell types including epithelial, immune, and mesenchymal cells. All quantitative data are presented as mean ± SD. Female mice are marked by unfilled legends.

ChP macrophages contribute to maintenance of epithelial tight junctions, which constitute the blood-CSF barrier. In 5xFAD mice, an increased proportion of macrophages contained intracellular Occludin (**Figure 5E**), suggesting enhanced tight junction remodeling. Consistent with barrier disruption, ChP epithelial junctions in 5xFAD mice displayed an irregular, wavy morphology compared to the linear pattern observed in WT controls (**Figure 5F**). We quantified this disruption using a linearity index derived from Occludin immunostaining on whole-mount explants, which revealed significantly disrupted linearity in 5xFAD mice (**Figure 5G**). Electron microscopy combined with intravenous labeling with horse radish peroxidase (HRP) corroborated these immunohistochemical findings (**Figure 5H**).

To test whether 5xFAD mice exhibit impaired barrier repair capacity, we challenged mice with intracerebroventricular (ICV) lipopolysaccharide (LPS). We previously demonstrated that ICV LPS causes severe disruption of tight junction proteins (including Occludin, Claudin-2, and ZO-1), with partial recovery by 5 days post-challenge^14^. As expected, both WT and 5xFAD mice displayed comparably disrupted tight junctions 24 hours following LPS while 5xFAD mice showed baseline damage consistent with what we observed before (**Figure S4**). However, 5xFAD females showed significantly delayed barrier recovery at 7 days post-challenge, assessed by Claudin-2 expression patterns and tight junction linearity index (**Figure 5J**). These findings indicate that amyloid-β pathology impairs the capacity of the ChP to restore barrier integrity following inflammatory challenge, especially in females.

ChP epithelial cells are long-lived^1^ and thus may be particularly vulnerable to accumulating senescence-associated damage. Given that recent work has shown that cellular senescence in the ChP drives age-dependent barrier dysfunction through irregular remodeling of tight junction proteins by senescent macrophages^58^, we asked whether amyloid-β pathology accelerates ChP cellular aging, We analyzed the ChP from *in situ* transcriptomics data generated from 9-month-old 5xFAD and WT mouse brains. Using a consensus senescence-associated gene signature^59-61^, we found that several ChP cell types in 5xFAD mice, including epithelial, mesenchymal, and immune cells, exhibited elevated senescence scores compared to WT controls (**Figure 5K-L**). These findings suggest that amyloid-β pathology promotes premature cellular aging in the ChP, which may contribute to our observations of dysregulated macrophage behaviors and barrier dysfunction in 5xFAD mice.

### Sexually dimorphic myeloid cell recruitment to the ChP in mouse AD models

Peripheral immune cell recruitment often accompanies barrier disruption. At 2 months of age, 5xFAD mice had increased numbers of ameboid CD45-high cells compared to WT controls (**Figure 6A-B**), consistent with increased peripheral cell infiltration. However, by 6 months, the difference was no longer significant (**Figure 6C**), paralleling the contrast we observed with macrophage coverage and process lengths (**Figure 4D-E**). These data suggest that peripheral immune cell infiltration is most pronounced during early pathology, at the onset of Aβ plaques in this model.

**Figure 6.**
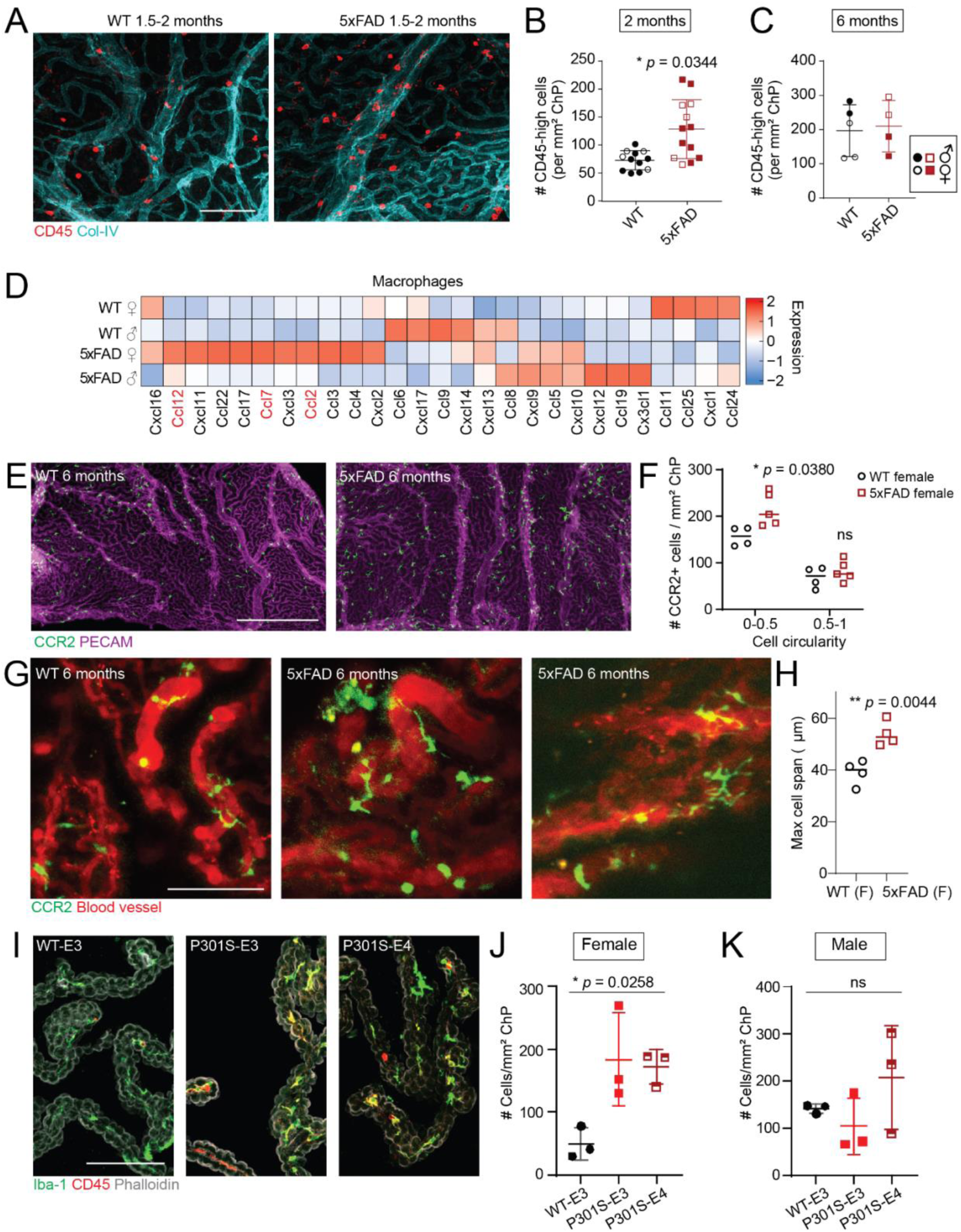
Myeloid cells are recruited to the ChP in mouse AD models with sex dimorphism. (**A**) Representative images showing increased CD45-high peripheral immune cells in the ChP of 5xFAD mice at 1.5-2 months old. Scale = 100 µm. (**B-C**) Quantifications showing increased CD45-high peripheral immune cells per mm^2^ ChP in only 1.5-2 months and not 6 months old 5xFAD mice. * p = 0.0344 (2 months), Welch’s t-test. (**D**) Gene expression of chemokines by sex and genotype from single-cell RNAseq of 5xFAD mice ChP at 6 months old; red = CCR2 ligand. (**E-F**) Representative images and quantifications showing female 5xFAD mice at 6 months old has increased number of CCR2+ cells with ramified macrophage-like morphology (low cell circularity). * p = 0.0380 (adjusted), Welch’s t-test with multiple comparison correction by Holm-Sidak method. (**G**) Representative snapshots of in vivo recording showing two examples of 5xFAD females at 6 months old having more macrophage-like CCR2+ cells. Scale = 100 µm. (**H**) Quantification of maximum span of individual CCR2+ cells from in vivo recordings showing female 5xFAD mice has increased cell span. ** *p* = 0.0044, Welch’s t-test. (**I-K**) Representative images and quantifications showing 9 months old female P301S mice had increased CD45-high Iba1+ macrophages, regardless of APOE genotypes. * *p* = 0.0258, Welch’s t-test. All quantitative data are presented as mean ± SD. Female mice are marked by unfilled legends unless separately marked.

Single-cell RNA-sequencing of the ChP from 6-month-old 5xFAD and WT mice revealed sex-biased transcriptional responses following the initial wave of immune cell infiltration ( **Figure S5, Supplemental Data 3**). Female 5xFAD macrophages specifically upregulated chemokines that recruit CCR2+ monocytes (e.g., *Ccl2*, *Ccl7*, and *Ccl12*) compared to female WT and male 5xFAD macrophages (**Figure 6D**). We also observed sex-dependent changes in the resident ChP macrophage pool. Using the *Ccr2^GFP^* reporter line to distinguish between recently recruited immune cells and ChP-resident macrophages, we classified GFP+ cells as ameboid (infiltrated monocytes) or ramified (newly differentiated macrophages). Female 5xFAD mice exhibited significantly increased numbers of GFP+ ramified cells compared to female WT controls (**Figure 6E-F**), indicating enhanced monocyte-to-macrophage differentiation. The difference was not observed in male mice due to high variations both at in WT and in 5xFAD mice. Using two-photon *in vivo* imaging, we captured a small number of CCR2+ cells within each FOV. CCR2+ cells in 5xFAD females spanned a greater distance than those in WT controls (**Figure 6G-H, Supplemental Video V4**).

To determine whether female-biased macrophage recruitment generalizes to other AD-relevant pathologies, we examined the P301S transgenic tau mouse model of tauopathy with a knock-in of humanized *APOE* ε3 or ε4 alleles at 9 months of age^62,63^. Consistent with our 5xFAD findings, we observed increased numbers of newly derived CD45-high macrophages in female, but not male, P301S mice compared to WT controls (**Figure 6I-K**). Male mice had higher level of CD45-high macrophages at baseline and did not show further recruitment in response to pathology. This macrophage recruitment was independent of human *APOE* allele status. Notably, P301S mice did not exhibit the increase in total or epiplexus macrophage numbers observed in 5xFAD mice, suggesting that epiplexus macrophage expansion may be specific to Aβ pathology, whereas enhanced monocyte recruitment and macrophage turnover at the ChP may represent a broader feature of AD-associated ChP inflammation.

## DISCUSSION

The ChP is the principal source of CSF and a critical barrier regulating molecular and cellular trafficking between the periphery and CNS, relaying inflammatory information between the brain and the periphery in both directions^14,22^. Despite mechanistic work demonstrating that inflammation in the ChP negatively impacts animal cognition^25^, clinical studies that increasingly link larger ChP volume to cognitive decline^37-39,64^, and CSF proteomics identifying ChP dysfunction as a distinct molecular subtype of AD^21^, the contribution of ChP pathology to aging and AD has not been systematically characterized. Here, we present a multi-modal atlas that was generated by integrating snRNA-seq, spatial profiling, AI-assisted quantitative histopathology, patient MRIs, and *in vivo* two-photon imaging paired with transcriptomics and mechanistic studies in mouse models to define the cellular and molecular landscape of ChP dysfunction in aging and AD.

Our work underscores the importance of a holistic and integrative approach to understand ChP pathology. Characterization of both human and mouse ChP at the transcriptional level provided valuable knowledge in understanding this brain barrier. Early work identified age-driven reduction in the expression of ion transporters and tight junction proteins by quantitative-PCR, hinting at reduced CSF production and barrier damage. ScRNAseq and snRNA-seq datasets, as well as epigenomics, from human tissues and animal models further revealed the molecular and cellular architecture of the ChP and its responses towards different brain conditions, such as healthy aging^28^, COVID infection^22^, and AD^42,43^. More recently, our *in vivo* imaging findings emphasize the importance of going beyond transcriptomics^14^. For example, we observed a diverse set of abnormal behaviors by ChP macrophages of 5xFAD mice (**Figures 4-6**), revealing complexity of ChP macrophage responses towards AD pathology that was not captured by transcriptomics. Therefore, a holistic approach that integrates unbiased transcriptomics datasets with studies of cell functions is essential to fully understand how the ChP functions in health and diseases.

A central finding of our study is that aging alone drives substantial ChP fibrosis, calcification, and macrophage dysfunction, whereas AD pathology exerts moderate additional effects with notable sex-differences. The broad age range of the TSDS study enabled us to establish that ChP structural changes emerge independent of dementia status, consistent with lifespan imaging studies showing ChP aging signatures appearing around age 30^65^ and transcriptomic shifts occurring later at 50-60 years old^66^. Furthermore, MRI studies demonstrate age-related increases in ChP volume are accompanied by reduced blood perfusion at the ChP^67,68^. This temporal framework wherein ChP pathology precedes the onset of clinical AD suggests that age-related barrier dysfunction may render the brain vulnerable to subsequent neurodegenerative insults. Altogether, our data support a model in which age-related ChP deterioration precedes and may facilitate AD progression. If so, interventions targeting ChP aging may represent a strategy to delay or prevent the development of AD, especially in high-risk populations. Distinguishing cellular changes driven by AD pathology versus normal aging is a major challenge for mechanistic studies of AD. Recent work suggest that the brain may take up two distinct trajectories of cellular changes, leading to either AD or alternative aging^69^. Further analyses of ChP fibrosis and calcification in patients with other age-related neurodegenerative disorders or in patients with familial AD or Down Syndrome who are at high risk of developing AD at relatively younger ages, may bring deeper understanding of the contribution of age-driven ChP fibrosis to neurodegeneration.

The stromal fibrosis and calcification we identified likely have significant functional consequences. Recent studies of meningeal lymphatics found that age-related ECM remodeling impedes CSF clearance^70^. Likewise, the increased fibrosis and calcification of the ChP stroma could block water movement from fenestrated capillaries to epithelial cells. Compounded by other age-associated changes in ChP epithelial cells including stunting of microvilli^71^, formation of Biondi bodies^35,36^, and downregulation of CSF secretory machinery (e.g., NKCC1^28,72^), the additional fibrotic changes in the ChP stroma may contribute to the well-established reduction of CSF secretion with advanced age^73^, and thereby allow amyloid-β buildup and the onset of Alzheimer’s disease. It is of further interest to understand the timeframe of the onset of different ChP pathologies and whether, in addition to cell autonomous changes, fibrosis and calcification could promote epithelial pathology by altering blood-to-epithelial cell trafficking.

Fibroblasts have emerged as key contributors to fibrosis and calcification following injury and disease in the central nervous system^74,75^, with recent studies showing that fibroblasts rapidly give rise to profibrotic myofibroblasts following CNS injury^75,76^. Critically, this response involves bidirectional crosstalk where macrophages promote myofibroblast activation through TGF-β, while activated fibroblasts reciprocally support macrophages through M-CSF, which is encoded by *CSF1*^75^. Moreover, fibroblast-mediated collagen deposition disrupts lymphatic drainage and CSF clearance^70^, which could negatively impact removal of Aβ or inflammatory cytokines in the milieu. Our snRNA-seq data revealed that inflammatory fibroblast subsets were enriched in AD (**Figure 1H**) and exhibited elevated expression of extracellular matrix factors (**Figure 3F**) as well as upregulated *CSF1* signaling **(Figure 3G**). Spatial transcriptomics demonstrated that fibroblasts and macrophages occupy overlapping stromal niches within the human ChP (**Figure 3A-E**), and cell-cell interaction analyses identified crosstalk between macrophages and inflammatory fibroblasts mediated by *CSF1* and TGF-β (**Figure 3F**), in line with mouse studies of CNS injury^75,76^. A concurrent multi-omics study of the ChP in the ROSMAP cohort similarly identified ECM pathway dysregulation in fibroblasts, but noted only mild collagen accumulation in AD, a discrepancy we attribute to the depth of our AI-assisted quantitative histopathology, which captured the strong age- and sex-driven effects on the development of ChP fibrosis^42^. Future studies may explore whether therapeutic targeting of fibroblast-macrophage interactions could limit the development of fibrosis and calcification in the ChP to preserve barrier function during brain aging, as suggested by studies in other brain compartments^77^.

Our mouse studies provide mechanistic insight into how macrophages contribute to barrier dysfunction during early AD-like pathology. *In vivo* two-photon live imaging of 5xFAD mice revealed that abnormal macrophage phenotypes, including increased phagocytic uptake of blood-sourced dextran, perivascular clustering, and reduced motility occurred prior to extensive plaque deposition and were accompanied by increased tight junction uptake in macrophages, irregular tight junction morphology, and delayed barrier repair upon inflammatory challenge. These findings extend previous work showing that ChP macrophages coordinate barrier disruption and repair through direct remodeling of epithelial tight junctions in acute inflammation^14^ and aging-associated senescence^58^. Analysis of spatial transcriptomics data from 5xFAD mice revealed elevated senescence signatures in ChP epithelial cells and macrophages, suggesting that amyloid pathology may promote premature senescence-associated dysfunctions in barrier maintenance (**Figure 5K-L**). Furthermore, barrier compromise can enable pathological immune cell entry into the CSF and brain parenchyma, where their abundance negatively correlates with cognition^10,11^. Our observation that infiltration of CD45-high cells during early-stage AD-like pathology suggests that macrophage-mediated barrier dysregulation may facilitate such infiltration. Future work should determine whether the molecular underpinnings of ChP macrophage dysfunction extend their influences beyond the blood-CSF barrier to affect brain-wide inflammation.

Sex differences in AD prevalence remain incompletely understood. MRI studies show that females exhibit a more than two-fold greater rate of ChP enlargement compared to males^38^, while mouse studies have shown female-biased hippocampal microglial senescence and inflammation during aging that may contribute to sex differences in AD prevalence^78^. We observed sex-dependent changes in both patient samples and mouse models. In human samples, we found that males had a stronger association between ChP fibrosis and calcification and AD pathology. In mouse models, we observed enhanced macrophage recruitment in the females of both the 5xFAD model for amyloid pathology and the P301S model for tauopathy. Our scRNA-seq of 5xFAD mice supported the histological observation by showing female-specific upregulation of macrophage-secreted CCR2 ligands. Recent work also reported sex differences in ChP macrophages and secretome, with immune signatures enriched in females^79^. Collectively, these data raise questions about whether males and females may have different mechanisms underlying ChP damage or repair, which then contribute to differences in AD risk and progression, warranting further investigation.

Our atlas highlights the value of combining postmortem human studies and early pathology investigations with animal models. However, our findings also reveal the limitations in the current models to study the ChP in aging and AD, shown primarily by the failure of mouse models to recapitulate pro-fibrotic transcriptomic signatures and pathology. Commonly used AD mouse models do not capture the complex genetic backgrounds of human patients. These models, especially 5xFAD mice, also have accelerated pathology and do not model age-driven changes like fibrosis. Other humanized models such as iPSC, microfluid 3D models, and organoids reflect the genetic complexity of humans and have demonstrated their potential in bridging human pathology studies and functional investigation *in vitro*^80-83^, but it remains challenging to retain cellular aging features as the process fosters rejuvenation. Likewise, common mouse models like 5xFAD mice have accelerated pathology, meaning the animals do not incorporate age-driven changes, such as fibrosis. New advances in human stem cell technology and humanized mouse models combining multiple known genetic contributors will provide more physiologically relevant AD models in the future.

## Limitations of the study

Our analysis on human pathology has several limitations: the TSDS cohort’s wide age range enabled identification of age-associated pathology, but the lack of detailed medical history makes it challenging to analyze AD associations compared to studies using other conventional AD-oriented brain banks. Second, the Vantaa85+ cohorts represent a more restricted age range (>85 years of age), and thus also limits our ability to disentangle disease phenotypes from age. Lastly, while we aim to delineate age contribution versus AD disease contribution to ChP pathology, in most patients, age and AD pathology are not independent. While our data demonstrate strong age-associated pathological changes in the ChP such as fibrosis, calcification, and macrophage types, the lack of a strong association with AD pathology after normalizing for age should not be interpreted as evidence that AD pathology has a weak impact on the ChP. In contrast, our studies in mouse models support a strong impact of AD pathology on ChP cell functions. In addition, our pathology specimens were restricted to portions of the lateral ventricle ChP. How ChP in the third and fourth ventricles change with aging and AD remains undetermined, and whether fibrosis and calcification demonstrate spatial specificity within each ChP is unknown. Future human studies with broader age ranges, more spatial coverage of ChP from different brain ventricles, and thorough disease diagnosis, including longitudinal imaging studies, will provide further understanding on ChP dysregulation in response to AD pathology.

## ACKNOWLEDGEMENTS

We thank members of the Lehtinen, Ordovas-Montanes, Myllykangas, Wyss-Coray, and Yang labs for helpful discussions, Andreea Luchian for her support with Aiforia methodology, Onerva Levälampi, Kristiina Nokelainen, Nina Hankonen for excellent technical assistance, the Boston Children’s Hospital Cell Discovery Network for advice on computational analysis, and Nancy Chamberlin for critical reading and editing of the manuscript. We thank the following facilities and personnel: Chinfei Chen, Hisashi Umemori, Cheng-Hao Chien, Harry Cramer, and the BCH IDDRC Cellular Imaging Core; Harvard Medical School Electron Microscopy Facility; FIMM Digital Microscopy and Molecular Pathology Unit supported by HiLIFE and Biocenter Finland for slide scanning services; and The Sanderson Center for Optical Experimentation at University of Massachusetts Chan Medical School. We thank the Vantaa 85+ Study establishers, researchers, and collaborators over the years as well as the participants, their families, and carers. We thank the personnel of Tampere University Histology Facility ‘Histocore’. We thank Lisa Keene, Emily Ragaglia, Aimee Schantz, and Kathryn Torrez, for incredible administrative support, John Campos and Mark Montine for data management, and Angela Wilson, Amanda Keene, Jenna Kelley, Julia Ryan, Flavia Ernau, Kim Howard, and Katie Miller for outstanding technical support. We are deeply grateful to the research participants and their families without whom this work would be impossible.

This work was supported by: BrightFocus postdoctoral fellowship in Alzheimer’s Disease research A2022026F and Boston Children’s OFD/BTREC/CTREC Faculty Career Development Award (H.X.); Howard Hughes Medical Institute James H. Gilliam Fellowship for Advanced Study (P.L.). The National Multiple Sclerosis Society (Postdoctoral Fellowship Grant, FG-2208-40289) (S.V.O.). Abramson Fund for Undergraduate Research, Stone Father-Daughter Fund for Undergraduate Research, and Herchel Smith Undergraduate Science Research Fellow (A.D.). BrightFocus Foundation A2022006F (V.D.L.), Alzheimer’s Association AARF-22-923219 (V.D.L.). ADRC Development Award (UW, P30AG066509 D.J.). The UW BioRepository and Integrated Neuropathology (BRaIN) Laboratory and Precision Neuropathology Core are supported by the National Institutes of Health (NIH) through the UW Alzheimer’s Disease Research Center (P30AG066509), the Adult Changes in Thought (ACT) study (U19AG066567 and R01AG060942), the BRAIN Initiative Cell Atlas Network (UM1 MH130981 and UM1MH134812), the Seattle Alzheimer’s Disease Brain Cell Atlas (U19AG060909), cooperative agreements (U24AG072458; U24NS133949; U24NS133945; U24NS135651; U01NS137500; and U01NS137484), the US Department of Defense (DoD W81XWH-21-S-TBIPH2), the Chan-Zuckerberg Initiative, and the Allen Institute for Brain Science. Dr. Keene is additionally supported as a Weill Neurohub Investigator and the Nancy and Buster Alvord Endowed Chair in Neuropathology. AHA–Allen Initiative in Brain Health and Cognitive Impairment (19PABHI34580007) (T.W.-C and A.C.Y.), the Michael J. Fox Foundation (T.W.-C.), The Phil and Penny Knight Initiative for Brain Resilience (T.W.-C. and A.I.). NIMH-R01MH113743 (D.P.S.), NINDS-R01NS117533 (D.P.S.), NIA-R01AG068281 (D.P.S.), Cure Alzheimer’s Fund (D.P.S.), the Miriam and Sheldon G. Adelson Medical Research Foundation (D.P.S.). U19AG069701 (D.M.H.), NS090934 (D.M.H.), and the Freedom Together Foundation (D.M.H.). This research was supported in part by the Intramural Research Program of the NIH (D.S.R.). The contributions of the NIH authors are considered Works of the United States Government. Cure Alzheimer’s Fund (D.S.R.); Adelson Family Foundation (D.S.R.); Jane and Aatos Erkko Foundation (P.J.K.); Finnish Foundation for Cardiovascular Research (P.J.K. and T.L.); Finnish Cultural Foundation (E.H.K.). National Institute of Neurological Disorders and Stroke 1R01NS128909-01 and 1RF1NS139975-01 (A.C.Y.). Academy of Finland (341007, L.M.), HUS Diagnostic Center and Helsinki University Hospital competitive research fund (TYH2022316, L.M.) Liv och Hälsa foundation (L.M.), Finska Läkaresällskapet (L.M) and Jane and Aatos Erkko Foundation (L.M.). The Pew Charitable Trusts Biomedical Scholars, The Broad Next Generation Award, NIH R01 HL162642, and The Cell Discovery Network at Boston Children’s Hospital supported by the Manton Foundation and Warren Alpert Foundation (J.O.-M.); The New York Stem Cell Foundation – Robertson Investigator (J.O.-M.); Cure Alzheimer’s Fund (M.K.L., and L.M.), Sigrid Jusélius Foundation (M.K.L.), and NIH R01 NS088566, NS129823, RF1048790 (M.K.L.); BCH IDDRC 1U54HD090255 and NIH grant #S10OD030322. The findings and conclusions presented in this paper are those of the authors and do not necessarily reflect the views of the NIH or the U.S. Department of Health and Human Services.

## DISCLOSURE

D.S.R. has received research funding from Abata and Sanofi. D.M.H. co-founded and is on the scientific advisory board of C2N Diagnostics. D.M.H. is on the scientific advisory boards of Denali, Genentech, Cajal Neuroscience, and Switch and consults for Pfizer, Novartis, and Roche. D.M.H. is on the advisory board of *Neuron* and *Cell*. J.O.-M. reports compensation for consulting services with Tessel Biosciences, Radera Biotherapeutics, and Passkey Therapeutics.

## AUTHOR CONTRIBUTIONS

H.X., P.L., D.J., A.C.Y., L.M., J.O.-M., and M.K.L. conceptualized and designed the study; H.X., P.L., B.E., J.O., J.W., R.P., D.K., S.V.O., V.D.L., D.J., D.P.S., E.H.K., D.S.R., A.C.Y., L.M., J.O.-M., and M.K.L. established methodology; H.X., P.L., B.E., L.I.J.B., J.W., K.C., H.P., A.P., J.B., M.E.G., V.D.L., E.H.K., A.C.Y., S.A., M.E.A, and D.J. conducted experiments; H.X., P.L., B.E., J.O., J.W., K.C., H.P.,M.I.M., J.T., J.F.H., S.A., S.V.O., A.T., A.D., K.E.P., J.F.H., V.D.L., D.S.R., A.C.Y., L.M., J.O.-M., and M.K.L. analyzed data; P.L. C.D.K., C.S.L., D.M.H., D.P.S., D.S.R., T.W.-C., A.I., P.J.K., T.L., and E.H.K. provided experimental material; D.J., D.S.R., T.W.-C., A.I., D.P.S., P.J.K., A.C.Y., L.M., J.O.-M., M.K.L. provided funding and supervised the study; H.X., P.L., B.E., J.O., J.W., L.M., J.O.-M., and M.K.L. wrote the manuscript. All co-authors read, revised, and approved the manuscript.

## METHODS

### Data and Code Availability

1. Human snRNA-seq data and mouse scRNA-seq data will be made publicly accessible on synapse.org (project number syn71304114) upon publication. The datasets are available for visualization through Broad Institute single cell portal under project ID SCP3449 (human) and SCP3451 (mouse).
2. Raw sequencing reads for mouse scRNA-seq data will be available in GEO upon publication. All other original data are available from the Dr. Lehtinen upon request. All biological materials were either directly commercially available or are available upon request.
3. All MATLAB code used in this study are published^55^ and available on Github at https://github.com/LehtinenLab/Shipley2020.
4. Availability of other source data and any additional information required to reanalyze the data reported in this paper are available from the lead contact upon request.

## EXPERIMENTAL MODEL AND SUBJECT DETAILS

### Human Subjects

#### Single-nucleus RNA-sequencing cohorts

Two independent cohorts of frozen lateral ventricle ChP were used. The first cohort was obtained from Rush University’s Religious Orders Study/Memory and Aging Project and the second was collected by the Neuropathology Core of the Alzheimer’s Disease Research Center (ADRC) of the University of Washington (UW) at autopsy. Tissue was flash frozen and stored at -80C. All samples were collected with approval from local ethics committees and in accordance with institutional review board guidelines. Complete neuropathological examination for all tissue samples was carried out as previously described.

#### Histopathology cohorts

1. The Vantaa 85+ study, a population-based cohort of the oldest old, consists of individuals aged >85 years living in Vantaa, Finland on April 1^st^, 1991 (n=601, n=282 analyzed). Their key demographic and neuropathologic variables are shown in **Supplemental Data 2**. Exact descriptions of the cohort have been published previously. Each participant and/or their relatives gave informed consent for the study. The collection of tissue samples and their research use was approved by the Finnish national authority of medicolegal affairs. The ethical committees of the Helsinki University Hospital and the City of Vantaa approved the study.
2. The Tampere Sudden Death Study (TSDS) is comprised of 515 white Caucasian men and 185 white Caucasian women aged 16–97 years subjected to a medicolegal autopsy at the Department of Forensic Medicine, University of Tampere, Finland, between 2010 and 2015. The TSDS series covered approximately 20% of deaths in the region during the timeframe of the study collection. According to Finnish legislation, the indications for a medicolegal autopsy include sudden unexpected out-of-hospital death, accident, suspected suicide, or homicide. In Finland, individual informed consent is not required from autopsy cases or their next of kin in a forensic study setting. The Ethics Committee of Pirkanmaa Hospital District (Permission numbers R09097 and R16199) and the National Supervisory Authority for Welfare and Health (Valvira Document number 564/05.01.00.06/2010) approved the TSDS protocol. Previous descriptions of the cohort have been reported^51-53^. A subset of cases (n=181) with a sufficient portion of ChP tissue attached to the hippocampus section were analyzed in this study. Key demographic and neuropathological variables are shown in **Supplemental Data 2**.
3. University of Washington cohort. As described above, choroid plexus tissue from the lateral ventricles was collected by the Neuropathology Core of the Alzheimer’s Disease Research Center (ADRC) of the University of Washington (UW) at autopsy (**Supplemental Data 2**). Paraffin embedded tissue blocks were sectioned, processed, and analyzed at University of Helsinki.

#### Spatial transcriptomics

Postmortem human brain tissue was obtained from the Lieber Institute (Johns Hopkins University) and processed by the Knight Initiative for Brain Resilience (Stanford University), with approval from local ethics committees.

#### Patient MRI

Clinical and imaging data was collected from participants with multiple sclerosis (MS) (including radiologically isolated syndrome, RIS)^84^ or MS-mimicking diseases under the institutional review board-approved National Institute of Neurological Disorders and Stroke protocol, “Evaluation of Progression in Multiple Sclerosis by Magnetic Resonance Imaging” (NCT00001248). Written, informed consent was acquired from participants. Participants were categorized as having relapsing MS (RMS) or progressive MS (PMS)^85^. Participants who did not meet diagnostic criteria for MS4 or RIS and demonstrated neuroinflammatory symptoms with abnormal MRI were categorized as other inflammatory neurological disease (OIND). Age at scan time, disease modifying therapy (DMT) status, Expanded Disability Status Scale (EDSS)^86^, Scripps Neurological Rating Scale (SNRS)^87^, Symbol Digit Modalities Test (SDMT)^88^, and Paced Auditory Serial Addition Test (PASAT)^89^ scores were recorded when available.

### Animal studies

The Boston Children’s Hospital IACUC and University of Massachusetts, Amherst approved all experiments involving mice in this study. The following genetic strains were obtained from JAX laboratory: C57BL/6J (000664), 5xFAD (034848), *Cx3cr1^GFP^*(005582), *Ccr2^GFP^* (027619), *Cx3cr1^CreER^* (020940), Ai14 (007914). 5xFAD mice (B6.Cg-Tg(APPSwFlLon,PSEN1*M146L*L286V)6799Vas/Mmjax, RRID:MMRRC_034848-JAX) was obtained from the Mutant Mouse Resource and Research Center (MMRRC) at The Jackson Laboratory, an NIH-funded strain repository, and was donated to the MMRRC by Robert Vassar, Ph.D., Northwestern University. All 5xFAD mice used in this study were bred by crossing 5xFAD male with C57BL/6J female, and the offsprings were housed separately by both genotype and sex. P301S mice were bred in the Holtzman lab and crossed with humanized APOE3 or APOE4^62,63^. Both male and female mice were equally included in the study. TTRmNeon mice were generated at Boston Children’s Hospital and published previously^14,90^. Animals were housed in a temperature-controlled room on a 12-hr light/12-hr dark cycle and had free access to food and water.

## METHOD DETAILS

### Isolation of nuclei from human choroid plexus tissue

#### ROSMAP cohort

Nuclei isolation was performed as previously described^22^. Briefly, 40 mg of choroid plexus tissue was thawed in 250 μl of 1% BSA with 0.2 U μl−1 RNase inhibitor. The tissue was incubated in 5 ml of lysis buffer (10 mM Tris, 10 mM NaCl, 3 mM MgCl2, 0.1% Nonidet P40 substitute (Roche/Sigma, 11754599001), 0.2 U μl−1 RNase inhibitor, and protease inhibitor) on ice for 10 min with gentle swirling every 2 min. 5 ml of 1% BSA was added and the tissue triturated 10 times with a 5-ml serological pipette. After centrifugation (500g, 5 min), pelleted nuclei were resuspended in 1% BSA with 0.2 U μl−1 RNase inhibitor, gently triturated 10 times with a 1-ml regular-bore pipette tip and filtered twice through a 70-μm and then a 40-μm strainer (Flowmi). Debris was inspected on a brightfield microscope and nuclei were counted on an TC20 automated cell counter (Bio-Rad) after the addition of Trypan blue.

#### University of Washington cohort

Frozen choroid plexus tissue (80-100 mg) was sampled on dry ice and mechanically minced with scalpels in sterile Petri dish on ice. Tissue dissociation method was adapted from a previous protocol^91^ with modifications. Tissue was transferred to 1.5 ml microcentrifuge tube (Fisherbrand ™ Pellet Pestle™ Microtubes 12-141-365) containing 250 µl TST buffer (Table 1-2). Tissue was manually homogenized with pestle (Fisherbrand™ RNase-Free Disposable Pellet Pestles, Cat. No. 12-141-364), pestle rinsed with 500 µl TST. Tissue was rotated at 4 °C for 10 minutes, and filtered through 40 µm filter (Corning, Cat. No. 352340). Samples were centrifuged at 500 *x* g, 4 °C for 5 minutes, and pellet was resuspended in 1 ml ST buffer with RNase inhibitor (Table 1) and DAPI staining solution (1/1000, Abcam, Cat. No. ab228549) for 10 minutes. Nuclei were isolated on the SONY cell sorter SH800S with a 100 µm chip and collected in nuclei suspension buffer (Table 3).

**Table 1:** 2X ST buffer.

| Chemical | Company | Cat. No. |
| --- | --- | --- |
| 292 mM NaCl | Thermo Fisher Scientific | AM9759 |
| 20 mM Tris-HCl pH 7.5 | Thermo Fisher Scientific | 15567027 |
| 2 mM CaCl<sub>2</sub> | VWR International Ltd | 97062-820 |
| 42 mM MgCl<sub>2</sub> | Sigma Aldrich | M1028 |

**Table 2:** TST buffer.

| Chemical | Company | Cat. No. |
| --- | --- | --- |
| 1 ml 2X ST buffer |  |  |
| 60 µl of 1% Tween-20 | Sigma-Aldrich | P-7949 |
| 10 µl of 2% bovine serum albumin (BSA) | Sigma-Aldrich | A3294 |
| 920 µl of nuclease-free water | Fisher Scientific | 10-977-023 |
| 10 μl (5 μl/mL) Protector RNase Inhibitor (2000 units) | Sigma-Aldrich | 3335399001 |

**Table 3:** Nuclei suspension buffer.

| Chemical | Company | Cat. No. |
| --- | --- | --- |
| 500 μl 10% bovine serum albumin (BSA) | Sigma-Aldrich | A3294 |
| 500 μl 1mM Aurintricarboxylic acid (ATA) in PBS | Sigma-Aldrich | A1895 |
| 4 ml 1X PBS | Fisher Scientific | AM9625 |
| (5 μl/ml) Protector RNase Inhibitor | Sigma-Aldrich | 3335399001 |

\*filtered prior to addition of RNase inhibitor

### Single-cell dissociation of mouse choroid plexus

The lateral ventricle (LV) ChP was dissected in cold RPMI media with 10mM HEPES and minced with a pair of fine scissors in a 1.5 ml tube (approximately 200 times until only small pieces remained). LV ChP from 3 adult male mice were pooled into each sample. Tissue pellets were digested in an enzyme cocktail containing collagenase P (0.5 mg/ml, Sigma cat. 11213865001), dispase (0.8 mg/ml, Worthington cat. LS02104), and DNAse1 (250 U/ml, Worthington cat. LK003172) for 30 min. at 37° C with slow head-to-toe rotation. Post-digestion tissues were washed, triturated, and filtered with 70 µm mesh. All pipette tips and tubes were coated by incubating in HBSS containing 2% BSA. For scRNA-seq preparation, to avoid transcriptional changes during the dissociation process, dissection and digestion solutions contained actinomycin (Sigma, cat. A1410, 5 µg/ml), triptolide (Sigma, cat. T3652, 10 µM), and Anisomycin (Sigma, cat. A9789, 27.1 µg/ml) ^92^. For CSF analysis, CSF from 5 adult male mice were collected and pooled into each sample. The CSF was treated with ACK lysing buffer (Thermo Fisher A1049201) to remove red blood cells. Cells from CSF were pelleted at 1000 g x 10 min. (4° C) and washed.

### Droplet-based sequencing

Human snRNA-seq:

Libraries were prepared using the Chromium Next GEM Single Cell 3ʹ v.3.1 according to the manufacturer’s protocol (10x Genomics), targeting 10,000 nuclei per sample. For the ROSMAP cohort, 15 cycles were applied to generate cDNA and all samples underwent 15 or 16 cycles for final library generation. Libraries were sequenced on a NovaSeq 6000 S4 Flow Cell. For UW cohort, the library was prepared following manufacturer’s instructions and sequenced by Northwest Genomics Services, University of Washington Genome Sciences on NovaSeq 6000 S2 Flow Cell.

#### Mouse scRNA-seq

Single cell suspensions were counted and adjusted to 1500 live cells per µl. Approximately 20,000 cells per sample were loaded onto the Chromium controller. Libraries were prepared according to manufacturer protocols (Chromium Next GEM Single Cell 3’ Reagent Kits v3.1 (Dual Index)). Sequencing was performed at Broad Institute on a NovaSeq 6000 SP Flow Cell.

### Single cell data processing

#### Human single-nucleus RNA-seq

FASTQs were demultiplexed and aligned to the human GRCh38 genome using CellRanger v6 (10X Genomics), counting reads mapping to pre-mRNA to account for unspliced nuclear transcripts. Three approaches were combined for quality control: (1) ambient RNA contamination was removed using SoupX^93^ for each individual sample; (2) outliers with a high ratio of mitochondrial (>10%, <200 features) relative to endogenous RNAs and homotypic doublets (>5,000 features) were removed in Seurat^94^; and (3) doublets were removed using DoubletFinder^95^. Nuclei with > 200 unique molecular identifiers were retained. The final dataset contained 197,108 nuclei.

Datasets from ROSMAP and UW cohorts were combined and batch-corrected using Harmony with default parameters. Cell type annotations were assigned based on canonical marker genes.

#### Mouse single-cell RNA-seq

Gene expression data were processed using CellRanger v8 with alignment to the mm39 reference genome followed by removal of ambient RNA contamination using CellBender v0.3.0^96^. We filtered cells with <300 detected features or >30% mitochondrial reads, yielding a total of 87,948 cells. Batch effects between experimental days were corrected using Harmony integration (lambda = 1).

Clustering was performed iteratively across at progressively refined resolutions to identify and remove doublets and low-quality cells. Label transfer from an annotated reference dataset was used to guide initial annotations^14^. Cell type annotations were assigned hierarchically at four levels (Class, Type, Subtype, State) based on canonical marker genes: epithelial (*Htr2c*, *Wdr17*, *Pcp4*, *Cd164*), fibroblast (*Coch*, *Dcn*), pericyte (*Notch3*, *Cox4i2*), smooth muscle cell (*Acta2*, *Myh11*, *Tagln*), endothelial (*Pecam1*, *Cldn5*), macrophage (*Csf1r*, *Pf4*, *Cd163*, *Lgals3*, *Cadm1*), microglia (*Siglech*), monocyte/DC (Ccr2), and lymphoid (*Cd3e*, *Cd8b1*, *Nkg7*, *Xcl1*).

#### Estimation of Variance

We performed variance decomposition using variancePartition^97^ on sample-wise average expression from the Harmony-integrated human object and fit linear mixed models with main effects for feature counts, mitochondrial percentage, study, APOE genotype, age of death, sex, Braak score, postmortem interval, Alzheimer’s disease status, and tissue processing batch, plus interaction terms (**Figure S1**).

Cell state taxonomy construction.

Hierarchical relationships among end-clusters in ChP cell types (cell “states”) were inferred using aggregated expression profiles, which were computed for each cell state across highly variable genes using a median absolute deviation (MAD)-based threshold on residuals from a loess-fitted mean-variance relationship. Expression profiles were z-scored across cell states, Euclidean distances were computed, and hierarchical clustering was performed using Ward’s metric (ward.D2).

#### Differential cell state abundance

Differential abundance of cell states between AD and control subjects was assessed using limma^98^ with arcsin-square root transformed proportions. Donors with fewer than 300 total cells were excluded to ensure robust proportion estimates. For each donor, the proportion of cells belonging to each cell state was calculated, then proportions were arc-square root transformed to stabilize variance and approximate normality. A linear model was fit with disease status with study of origin (ROSMAP or UW) as a covariate: transformed proportion ∼ AD_status + study. Empirical Bayes moderation was applied using robust estimation. Log odds ratio were calculated to quantify effect size, with positive values indicating enrichment in AD relative to Control, and p-values were adjusted for multiple testing using the Benjamini-Hochberg FDR method.

#### Pseudobulk differential gene expression and gene set enrichment analysis

Differential gene expression between AD and Control was assessed using DESeq2^99^ on pseudobulk expression profiles, where raw counts were aggregated by summing across all cells of a particular cell type belonging to each donor. Counts matrices were filtered to retain genes detected in at least 30% of samples with expression in at least 5% of cells within the population. The DESeq2 object was generated with the design formula: ∼ AD_status + Age_scaled + Sex. Gene set enrichment analysis (GSEA) was performed using fgsea^100^ on DESeq2 results using the Hallmark gene sets from msigdbr. Pathways with adjusted p-values < 0.05 were considered significant.

#### Cell-Cell Communication Analysis

Intercellular communication was inferred using MultiNicheNet^47^, which integrates differential expression analysis with *a priori* knowledge of ligand-receptor and ligand-target relationships. First, we excluded neural and glial cells (including microglia) and filtered cell types with fewer than 10 cells per sample and present in fewer than 30% of samples within a disease group. Next, we filtered genes that were detected in fewer than 5% of cells and fewer than 30% of samples per condition. We then generated pseudobulk expression profiles per cell type per sample for differential gene expression. For AD vs. Control differential interaction analysis (Figure 3), we used the contrast AD – Control and included study of origin as a covariate. For analysis of differential ligand activity between AD, MCI, and Control, we set the contrast to AD – (Control+MCI)/2, MCI – (AD+Control)/2, Control – (AD+MCI)/2. Genes with log2FC and p<0.05 were considered significant.

#### Ligand activity inference

For each receiver cell type, ligand activities were inferred based on the enrichment of ligand target genes among differentially expressed genes. The NicheNet activity score reflects the regulatory potential of each ligand to induce the observed transcriptional differences. The top 250 target genes per ligand were considered for activity scoring and only upregulated ligand activities were retained.

### Pathological analysis of human choroid plexus

#### Pathological procedures

1. Vantaa 85+ cohort: During the course of the study, brains harvested at autopsy were fixed for at least two weeks in formalin before sampling. Neuropathological examination was performed for all cases as described before^48-50^. Specimens of the right lateral ventricle ChP embedded in paraffin were used for this study.
2. TSDS cohort: Individual tissue samples (cerebellar cortex, frontal cortex, hippocampus, insula-putamen, pons, and substantia nigra) from unfixed brains were collected at forensic autopsy and stored in phosphate-buffered 4% formaldehyde solution for at least 48 hours prior to paraffin embedding into TissueTek cartridges^51-53^.
3. UW cohort: UW AD Research Center (ADRC) Neuropathology Core handled tissue evaluation and processing blind to clinical diagnosis as previously described^101-103^. Brain tissues used in this study were immersion-fixed in 10% neutral buffered formalin for at least 2 weeks and evaluated (wholly and after coronal sectioning) for any gross lesions. Tissue samples were dissected, processed, and embedded in paraffin prior to sectioning and staining.

#### Anti-Iba1 immunohistochemistry

4 µm thick paraffin sections were mounted on glass slides and dried overnight at 37°C, followed by incubation at 56°C for 30 min. - 1h. After automated deparaffinization (TissueTek DRS, Sakura Finetek Europe B.V., Alphen aan den Rijn, the Netherlands) heat induced antigen retrieval was performed at pH 6, for 20 min at 95 °C (Dako Target retrieval solution K2005, Agilent, Santa Clara, CA, USA), followed by formic acid treatment for 5 mins (Fisher Chemical Formic acid 98/100% Code: F/1900/PB15, Thermo Fisher Scientific, Waltham, MA, USA). Anti-Iba1 immunostaining (Wako polyclonal rabbit anti Iba1, #019-19741, FUJIFILM Biosciences, Santa Ana, CA, USA; diluted at 1:1500 for 30 mins ([Dako AB diluent S2022, Agilent])), was carried out on a Labvision Autostainer 480 (Thermo Fisher Scientific) using the Dako EnVision FLEX –kit (K8002, Agilent). Counterstain was performed with Mayer’s hematoxylin for 1-2 min.

#### Masson’s Trichrome staining

Tissue sections (4 µm) were mounted on glass slides and dried overnight at 37 °C. The next day, slides were incubated at 56 °C for 30 min before staining. Automated deparaffinization and hydration (TissueTek DRS, Sakura Finetek Europe B.V., Alphen aan den Rijn, the Netherlands) were performed prior to overnight fixation in Bouin’s solution (HT10132, Sigma-Aldrich, Saint Louis, USA) at RT followed by rinsing with tap water until clear. The slides were stained with Weigert’s Iron Hematoxylin (HT1079, Sigma-Aldrich, Saint Louis, USA) for 5 min after which they were placed under running tap water for 5 min and rinsed with Aqua. The samples were then stained with Biebrich Scarlet-Acid Fuchsin solution (HT151, Sigma-Aldrich, Saint Louis, USA) for 5 min following aqua rinse 3 x 1 min. Next, the slides were incubated in Phosphotungstic acid solution (HT152, Sigma-Aldrich, Saint Louis, USA) and Phosphomolybdic acid solution (HT153, Sigma-Aldrich, Saint Louis, USA) in aqua (1:1:2) for 5 min following staining with Aniline Blue solution (B8563, Sigma-Aldrich, Saint Louis, USA) for 5 min. To remove excess dye, the sections were treated with 1 % acetic acid for 2 min followed by aqua rinse 3 x 1 min. The sections were then dehydrated through a series of alcohols and cleared with xylene before applying coverslips. Slide scanning was performed at FIMM Digital Microscopy and Molecular Pathology Unit, Helsinki, Finland, on a Panoramic 250 Flash III (3D HISTECH, Budapest, Hungary).

#### Micro-CT

Samples (n=253 from Vantaa 85+ cohort) were scanned using a Quantum GX microCT scanner (PerkinElmer, MA USA) with a peak voltage of 90 kV and a current of 88 µA. The imaging was done in a 36 mm FOV with Cu 0.06+Al 0.5 X-ray filter and with ring reduction. High-resolution scan mode was used with an exposure time of 4 min and a voxel size of 72 µm. The images were captured over a 360° rotation around the sample. Tomograms were reconstructed with 3D Slicer (version 5.6.1) software. ChP tissue was manually segmented from the cassette and surrounding hippocampal tissue for the calcification volume analysis. Paraffin (and paraffin-infiltrated soft tissue) was excluded from the volume analysis by setting a threshold. The mean Hounsfield Unit (HU) value for paraffin was calculated from 10 different paraffin blocks that were manually thresholded. The same threshold (-16 HU) was used for all samples within the cohort. An estimation of the amount of calcification in each sample was calculated by dividing the acquired calcification volume (mm^3^) by an estimated tissue volume (mm^3^). An estimate of the tissue volume was calculated from the tissue area (Aiforia) and thickness of the tissue (3D slicer). Samples that failed tissue recognition in Aiforia or had too little material left in the paraffin block were excluded from the Micro-CT analysis.

#### Neuropathologic evaluation

For the Vantaa 85+ study population, we used previously described Thal β-Amyloid phase, Braak NFT stage^104^ and Consortium to Establish a Registry for Alzheimer’s Disease (CERAD) neuritic plaque score^48^ data.

#### AIFORIA analysis

We created an artificial intelligence model using Aiforia® Create version 6.4. (Aiforia Technologies Oyj, Helsinki, Finland). The development of the convolutional neural network (CNN) algorithm started by selecting 38 cases for training. Two training layers were created: ChP tissue with area detection (parental layer) and another layer with object detection of villi (normal and fibrotic) and psammoma bodies (child layers). The normal villi category included villi that had a typical structure; epithelial cells lining the villus and a visible capillary, any villi with dense fibrosis and missing capillary were annotated as fibrotic^34^. The psammoma body category included calcified lamellar structures that often appeared in groups^105^. To set the ground truth, a total of 571 training regions (305 for ChP and 266 for the object classes) including background and artifacts were annotated. The total number of object annotations was 6902 (2145 for psammoma bodies, 2926 for normal villi and 1831 for fibrotic villi). Training parameters are shown in **Supplemental Data 2**. The training resulted in a model with a total area error of 0.39 % (F1 score: 97.81 %) for the ChP tissue and a total object error of 5.64 % (F1 score: 97.15 %) for the object classes. A visual pre-validation was carried out for a small set of cases from the TSDS, the Vantaa 85+, and the UW cohorts by the annotator. The Aiforia validation tool was used for the final validation by three experienced neuropathologists (L.M., B.E., H.P.). For validation, four cases that had not been used for training were selected. The external validators performed 18 area annotations for the ChP tissue and 98 object annotations for the object classes on selected areas. Among the three external validators, error for the ChP tissue was 6.52 % (F1 score: 91,43 %) and for the object classes (villi and psammoma bodies) 0.67 % (F1 score: 99,66 %). Quantification of the villi (normal and fibrotic) and psammoma bodies was carried out by image analysis with the algorithm. The analysis also determined the area (mm^2^) for ChP tissue in each sample. The amount of fibrosis was calculated by dividing the number of fibrotic villi by the number of normal villi and the number of psammoma bodies was divided by the tissue area. After running the image analysis, manual quality check was performed on each sample. Samples that had poor tissue recognition due to defective material or under 45 normal villi detected by the algorithm were excluded from the analysis.

To analyze macrophages, we developed another CNN AI model for image analysis, employing Aiforia® Create. At first, 50 cases were carefully selected for training. Specimens obtained from the Helsinki Biobank were included to facilitate AI training on a heterogenous and representative sample set. Segmentation was performed using two distinct training layers within the training pipeline: A region layer for detection of the ChP and another region layer covering different classes of macrophages (amoeboid, dystrophic, reactive, rod-shaped).

For training the algorithm, ChP macrophages were defined in resemblance to previously published morphological descriptions of central nervous system microglia subtypes^106,107^. Representative micrographs of ChP macrophages are depicted in **Figure 3**.

1. Amoeboid macrophages show a rounded or oval, usually well delineated cell body with smooth edges and no or only few, very short and stubby protrusions.
2. Dystrophic macrophages: Irregular cell body, with beaded/fragmented processes and deteriorated cytoplasm. Borders might be hard to delineate.
3. Reactive macrophages: These show an enlarged cell body with marked, thickened protrusions.
4. Rod-shaped macrophages: Thin, elongated rod- or club-shaped cell body, often found in perivascular location, sometimes also appearing curvilinear, comma- or crescent-shaped.

A total of 774 training regions (278 for ChP and 496 for macrophage layers), including tissue background and artifacts, were annotated to set a ground truth. Training parameters can be found in **Supplemental Data 2**. Verification of the algorithm revealed a total area error for the ChP layer of 0.01% (F1 score: 97.32%) and a total area error of 3.52% (F1 score: 89.12%) for the macrophage layers. After yielding sufficient sensitivity and precision, validation of the algorithm was carried out by i) visual pre-validation on a small subset of region of interest (ROI) by the annotator and ii) using the Aiforia validation tool. Validation using Aiforia was performed by three experienced pathologists (L.M., M.I.M., H.P.). For validation, eight cases with 51 regions (16 ChP, 35 macrophages), not previously used for training were selected. Average total area errors among the three validators were: 0.13% (ChP), 9.38% (macrophages), average F1 scores: 84.66% (ChP), 81.69% (macrophages).

For eventual quantification of ChP macrophages, a representative ChP ROI was marked on scanned micrographs of anti-Iba1 immunostained slides. ROIs were selected so that most of the ChP area with preserved histological architecture was included, whenever possible. As a readout of the image analysis run, the ChP area as well as the ChP macrophage cell count and % (area of macrophage class per total ChP area, termed “coverage” as of now) were determined by the algorithm in Aiforia. A fraction of macrophages per sample (total sum of each macrophage class divided by the total number of macrophages, in %) was calculated, in addition to the coverage used in subsequent statistical analyses. Before and after running the algorithm, every micrograph underwent a thorough quality check, particularly focusing on staining- and scanning quality and assessing if the algorithm performed sufficiently, yielding 230 Vantaa 85+ and 59 UW samples for further analysis.

### MRI scans of patients

#### Image acquisition and analysis

20 7T T2*-gradient weighted echo (T2* GRE, TR/TE1/TE2/TE3/TE4/TI = 72.7/12.6/23.75/34.8/46.05/15ms, flip angle = 12°, 0.5mm isotropic) and 7T T1-weighted magnetization prepared 2 rapid gradient echo (MP2RAGE, TR/TE/TI1/TI2 = 6,000/3.02/800/2,700ms, flip angle = 4/5°, 0.7mm isotropic) images were acquired on a Siemens Magnetom scanner.

Quantitative susceptibility images were generated from 7T T2* GRE magnitude and phase images. A brain mask was generated from the magnitude images using SynthStrip and eroded by one voxel. Total-generalized-variation (TGV) based method for QSM reconstruction^108^ (available at: https://github.com/QSMxT/TGV_QSM) was applied to the phase image and corresponding eroded brain mask for each echo. The four resulting QSM images were averaged with 3DCalc in AFNI^109^.

ChP was segmented on 7T T2* GRE magnitude images using 3D-Slicer (https://www.slicer.org/)^110^. Segmentations were created in a semi-automated fashion by AAT by applying the voxel thresholding tool dynamically to adequately segment the lateral ChP after training (by S.V.O., neurologist with 7 years of experience in neuroimaging). Segmentations of calcified regions were created by manually segmenting hypointense regions within the choroid plexus on 7T T2* QSM **(Figure 2F)**. ChP and calcification volumes were recorded. The calcification ratio (calcification volume divided by ChP volume) was calculated. The mean ( ±SD) of QSM values of the ChP segmentations were recorded. The automated segmentation tool Pseudo-Label Assisted nnU-Net (PLAn)^111^ was applied to 7T T1-MP2RAGE images to identify CSF, gray matter (GM), white matter (WM), and white matter lesions (WML). Fractional GM (GM-F) and fractional CSF (CSF-F) were calculated by dividing GM and CSF volumes, respectively, by normalization to the total intracranial volume.

#### Statistics

Correlations were assessed with a Spearman correlation. Unpaired comparisons across categories were assessed with a Mann-Whitney U test. To account for age, a multiple linear regression model was performed.

#### Cohort characteristics

|  |  |
| --- | --- |
| Age (Mean ± SD) | 47.35 ± 12.72 (n=20) |
| Sex (F/M) | 16/4 |
| Diagnoses (MS/OIND) | 15/5 |
| MS Subcategorization (RMS/PMS) | 11/4 |
| Treatment Status (On/Off or Naïve) | 10/10 |
| EDSS (Median, IQR) | 3.0, 1.5-3.5 (n=15) |
| Scripps (Mean ± SD) | 88.1 ± 9.8 (n=19) |
| PASAT (Mean ± SD) | 54.2 ± 6.4 (n=11) |
| SDMT (Mean ± SD) | 53.9 ± 10.2 (n=15) |
| GM-F (Mean ± SD) | 0.458 ± 0.032(n=20) |
| WML (ml) (Mean ± SD) | 3.88 ± 4.13 (n=20) |
| CSF-F (Mean ± SD) | 0.193 ± 0.041 (n=20) |
| Calcification Ratio (Mean ± SD) | 0.114 ± 0.077 (n=20) |
| CP Volume (ml) (Mean ± SD) | 1.491 ± 0.452 (n=20) |
| CP QSM Susceptibility (ppm) (Mean ± SD) | -0.00120 ± 0.00793 (n=20) |
| Calcification Volume (ml) (Mean ± SD) | 0.180 ± 0.153 (n=20) |

### Spatial transcriptomics of human ChP

#### Xenium Sample Processing

Postmortem human brain tissue was obtained from the Lieber Institute (Johns Hopkins University) and processed by the Knight Initiative for Brain Resilience (Stanford University), with approval from local ethics committees. Fresh frozen tissue sections were obtained from the lateral ventricle of a cognitively normal 48-year-old male donor. Sections were cut at 10 µm thickness using a cryostat and mounted on Xenium-compatible slides (10X Genomics). Slides were stored at −80 °C until processing.

Xenium slides were processed according to the “Xenium *In Situ* for Fresh Frozen Tissues-Fixation & Permeabilization” demonstrated protocol (10x Genomics, CG000581), beginning with the 20-minute fixation step. Following permeabilization and cassette assembly, slides were processed according to the “Xenium *In Situ* Gene Expression with Cell Segmentation Staining User Guide” (10X Genomics, CG000749). Cell segmentation enables high–resolution cell boundary mapping. After hands-on slide processing, slides were loaded onto the Xenium Analyzer and processed.

#### Spatial Transcriptomics Alignment, Quality Control, and Annotation

A morphology stain (H&E) was applied to slides after the 10X Xenium workflow to assist with downstream spatial alignment and analysis. Images were processed and registered using Xenium Explorer v3.0.0, applying default autofluorescence correction and alignment parameters. Cells were retained for downstream analysis if they exhibited expression of at least ten transcripts distributed across a minimum of three distinct genes. Genes were retained if they were detected in greater than three cells. These filters were implemented in Scanpy (v.1.10.4)^112^.

Dimensionality reduction was conducted using scVI (v.1.2.0)^113^, with two layers and 30-dimensional latent representation, applied to the integrated dataset. A categorical batch covariate was included during training to account for inter-run variability and to mitigate batch effects. The resulting latent representations underwent Leiden clustering. Cell type annotations were transferred from our human single-cell dataset via logistic regression using shared features.

#### Spatial Neighborhood and Niche Analysis

Computational inference of spatial niches was performed using an Adversarially Regularized Graph Variational Autoencoder (ARGVA)^114^. The encoder consisted of a two-layer Graph Convolutional Network (GCN) that takes gene expressions and adjacency matrices (constructed from cellular neighborhoods within a 100-unit radius) as inputs, then transforms them into latent representations, producing both point estimates and uncertainty values. The gene expression input consisted of log-normalized transcript counts restricted to the predefined Xenium gene panel.

For decoding, we used an inner-product decoder for edge prediction to estimate the probability of two cells being within the same neighborhood^115^. To regularize the latent space and improve robustness, a discriminator neural network was used in an adversarial training framework, trained via logistic loss (a binary classification task). This adversarial regularization empirically improves both reconstruction fidelity and downstream clustering performance^114,115^. The objective function of our ARGVA combines (a) network reconstruction loss, (b) discriminator-derived regularization for noise-robust latent distributions, and (c) KL divergence to encourage alignment between the learned and prior distributions^116^. Model training was conducted over 10,000 epochs. Leiden clustering was performed on the derived latent space using a resolution parameter of 0.05. Analysis for this section was performed using PyTorch (v.2.4.0) and PyTorch Geometric (v.2.5.3).

The spatial colocalization of macrophages with other cell types was assessed by identifying the nearest cell type neighbor for each cell and calculating the relative frequencies of such proximities. These frequencies were normalized within each computationally defined niche to facilitate comparative analysis. This analysis was performed using custom scripts in Python (v.3.12.1).

### Mouse tissue collection and processing: explants and brain blocks

Animals were anesthetized by ketamine, perfused with ice cold PBS, and then followed by cold 4% paraformaldehyde (PFA). For cryosectioning, brains were dissected, further fixed in 4% PFA at 4° C overnight, and then incubated in 30% sucrose for at least 48 hours, followed by OCT (1 hour on ice) prior to freezing by dry ice and 2-met-butane bath. For wholemount ChP explant, the LV ChPs were dissected and incubated in 4% PFA at room temperature for 5 min., and then immunostained.

### Immunostaining of mouse samples

Cryosections and explants were blocked and permeabilized (0.3% Triton-X-100 in PBS; 5% serum), incubated in primary antibodies overnight and secondary antibodies for 2 hours. Sections and explants were counterstained with Hoechst 33342 (Invitrogen H3570, 1:10,000) and mounted using Fluoromount-G (SouthernBiotech).

The following primary antibodies were used: chicken anti-GFP (Abcam ab13970; 1:1000), rat anti-PECAM (BD Pharmingen 550274, 1:100), rat anti-CD45 (Fisher Scientific BDB550539, 1:50), rabbit anti-Iba1 (Wako 019-19741, 1:200), goat Anti-Type IV Collagen-Alexa Fluor® 488 (Southern Biotech 1340-30, 1:200), rabbit anti-Occludin (ThermoFisher 71-1500, 1:50), rabbit anti-Claudin 2 (ThermoFisher 51-6100, 1:100) Secondary antibodies were selected from the Alexa series (Invitrogen, 1:500).

### Confocal microscopy and image analysis

All confocal images were acquired using Zeiss LSM 880 and Zeiss LSM 980 with Airyscan confocal microscope. Explant images were taken with 2.5 µm z-stacks. All images were processed with FIJI^117^.

### Headpost and cranial window placement

Mice used for *in vivo* two-photon imaging (4-6 months) were outfitted with a headpost and a 3 mm cranial window over the left lateral ventricle at least three weeks prior to imaging as previously described^14,55^.

### Two-photon imaging and image processing

Two-photon microscopy (Olympus FVMPE-RS two-photon microscope; 512 x 512 pixels / frame) was used to record immune cells activities in ChP *in vivo* from 5xFAD mice or their WT littermate controls crossed with *Cx3cr1^GFP^* (JAX 005582) mice or *Ccr2^GFP^* (JAX 027619) mice. 200 µm Z-stack was acquired with 5 µm stepping size at each field of view over the course of approximately 1 hour. Imaging and analyses were performed as described^14,55^. A 25X magnification, 8 mm working distance objective was used. Images were registered and processed following the established algorithm^14,55^.

### Linearity index analysis of Occludin and Claudin 2

Individual cell borders were isolated in FIJI. The total length of the border was measured by manual tracing. The shortest distance was measured by drawing a straight line connecting the two ends of the border. Linearity index was calculated as Total length / straight line length

### Intracerebroventricular injection (ICV) of lipopolysaccharides (LPS)

Adult mice were anesthetized with isoflurane from a vaporizer. Following a midline scalp incision to expose the skull, the lateral ventricle was located by stereotaxic coordinates (bregma -0.4 mm, midline +1.0 mm, dura -2.2 mm). A single delivery path was prepared above the lateral ventricle. 5 µl of LPS solution (Sigma L3024, 0.5 mg/ml) was delivered using a Hamilton syringe into the lateral ventricle over the course of 5 min.

### HRP injection and TEM

As previously described^90^, mice were weighed and injected retro-orbital under anesthesia (isoflurane vaporizer) with 0.5 mg horseradish peroxidase (HRP Sigma P8250-200 KU) per g bodyweight diluted in sterile PBS. HRP was diluted such that each mouse received 100uL per 25 g bodyweight. HRP was allowed to circulate for 8 min, and then brains were harvested, frontal pole was removed, and fixed 1 hour in Karnovsky’s fixative (5% glutaraldehyde, 4% PFA, 0.4% CaCl2, in 0.1 M cacodylate buffer - Electron Microscopy Sciences), then fixed in 4% PFA in 0.1 M cacodylate at 4 °C overnight on a shaker. After washing the brains thoroughly in 20 mM glycine followed by ice cold 0.1 M cacodylate buffer, the lateral ventricle ChP were dissected and processed for DAB (Millipore Sigma D5905). DAB solution was made using 5 mg DAB in 9 mL of 0.1 M cacodylate plus ∼9 mM H2O2. The ChP were developed at room temperature until dark brown (1-2 minutes). The ChP was then processed, sectioned, and imaged at the Conventional Electron Microscopy Facility at Harvard Medical School. Tissue was postfixed with 1% osmiumtetroxide (OsO4)/1.5% potassium ferrocyanide (KFeCN6) for one hour, washed in water three times and incubated in 1% aqueous uranyl acetate for one hour. This was followed by two washes in water and subsequent dehydration in grades of alcohol (10 min each; 50%, 70%, 90%, 2 × 10 min 100%). Samples were then incubated in propyleneoxide for one hour and infiltrated overnight in a 1:1 mixture of propyleneoxide and TAAB Epon (Marivac Canada Inc. St. Laurent, Canada). The following day, the samples were embedded in TAAB Epon and polymerized at 60 degrees C for 48 h. Ultrathin sections (about 80 nm) were cut on a Reichert Ultracut-S microtome, and picked up onto copper grids stained with lead citrate. Sections were examined in a JEOL 1200EX Transmission electron microscope or a TecnaiG² Spirit BioTWIN. Images were recorded with an AMT 2k CCD camera.

### Multiplexed error-robust in situ hybridization (MERFISH): sample preparation and imaging

Mice were anesthetized and euthanized by decapitation. Brains were extracted and immediately transferred to cold OCT. Embedded brains were frozen and stored at -80°C. Vizgen’s fresh frozen tissue sample preparation protocol was followed. Briefly, brains sections were sliced at 10 μm. Tissue slices were mounted on Vizgen’s bead-coated functionalized coverslip. Once adhered to the coverslip, the samples were fixed (4% PFA in 1× PBS, 15 min, room temperature) and washed (3x, 5 min, 1X PBS). Then samples were incubated overnight in 70% EtOH to permeabilize the tissue. Following permeabilization samples were incubated for 30 min in Formamide Wash Buffer (30% deionized formamide (Sigma, S4117) in 2× SSC (Thermo Fisher, AM9765)) followed by the addition of the gene library mix for the hybridization step (48h, 37 °C). Samples were then washed twice with Formamide Wash Buffer (2x, 30 min, 47°C), the tissue was embedded in a gel embedding solution (0.5% of 10% w/v ammonium persulfate, 0.05% TEMED, 4% 19:1 acrylamide/bis-acrylamide solution, 0.3M NaCl, 0.06M Tris PH8) followed by overnight incubation with tissue clearing solution (2X SSC, 2% SDS, 0.5% v/v Triton X-100, and proteinase K 1:100) at 37°C. Finally, tissue was washed, incubated with DAPI and polyT solution (15 min RT), and washed with formamide wash buffer (10 min RT). After sample preparation, the MERSCOPE 500 gene imaging kit was activated by adding 250 μl of Imaging Buffer Activator (VIZGEN, #203000015) and 100 μl of RNAse Inhibitor (New England BioLabs, M0314L). 15 ml of mineral oil was overlaid on top of the imaging buffer through the activation port and the instrument was primed and the imaging chamber was assembled as per MERSCOPE user guide. A 10x low resolution mosaic of the sample was then acquired, the imaging area was selected, and the sample was imaged.

### MERFISH post-i2maging data processing and analysis

Following image acquisition, the resulting data were decoded using Vizgen’s analyzing pipeline incorporated in the MERSCOPE. Vizgen’s postprocessing tool (Vizgen, Cambridge, MA) was then applied to obtain the cell segmentation based on the DAPI staining by using the CellPose algorithm. Segmentation was performed on the middle Z plane (3rd out of 7) and cell borders were propagated to z-planes above and below. MERFISH processed data were analyzed in RStudio using Seurat 4.1.3, R 4.2.2 and custom-made scripts. Cell filtering was applied to the dataset to remove cells <100 µm^3^ and <10 unique transcripts and <40 transcript counts. Cell gene expression per cell was then normalized to each cell’s volume and the mean RNA per sample. Following annotation of choroid plexus (ChP) by region, ChP cells were subclustered and annotated by cell type using known marker genes. To assess differences in senescence gene expression across genotypes and cell types, we subset ChP cells by major cell type (Epithelial, Mesenchymal, Endothelial, & Immune) and compared senescence scores between wild-type (WT) and 5xFAD animals using the Wilcoxon rank-sum test. Senescence is evaluated by a set of genes from the recently published SenMayo^59^ signature. Effect sizes were calculated using Cohen’s d (effsize v0.8.1). P-values were adjusted across cell types using the Benjamini-Hochberg method to control the false discovery rate. Visualization of the SenMayo gene modules in WT and 5xFAD ChP was performed on the Vizgen MERSCOPE Visualizer software.

### Computational analysis, statistics and schematics

All single-cell RNA-seq analyses were performed in R (v4.5.1). Single-cell data processing utilized Seurat (v5.3.1)^94^ with harmony (v1.2.4)^45^ for batch correction. Differential cell type abundance was assessed using speckle (v1.10.0)^118^ with limma (v3.66.0)^98^, adjusting for study as a covariate. Pseudobulk differential expression analysis employed DESeq2 (v1.50.2)^99^. Pathway enrichment analysis used fgsea (v1.36.0)^100^ with gene sets from MSigDB (msigdbr v25.1.1), and gene signature scoring was performed using AUCell (v1.32.0)^119^. Cell-cell communication analysis employed multinichenetr (v2.1.0)^47^. Figures were generated using ComplexHeatmap (v2.26.0)^120^ and circlize (v0.4.17)^121^ for circos plots. Two-photon imaging data were processed with MATLAB pipelines as published and available on GitHub^14,55^. Statistical tests are specified in the respective sections in Methods. Schematic diagrams were created in parts with BioRender.com. Xenium visualization was performed using Matplotlib (v3.9.2) and Seaborn (v0.13.2)^122^.

## SUPPLEMENTAL FIGURES AND FIGURE LEGENDS

**Supplemental Figure 1.**
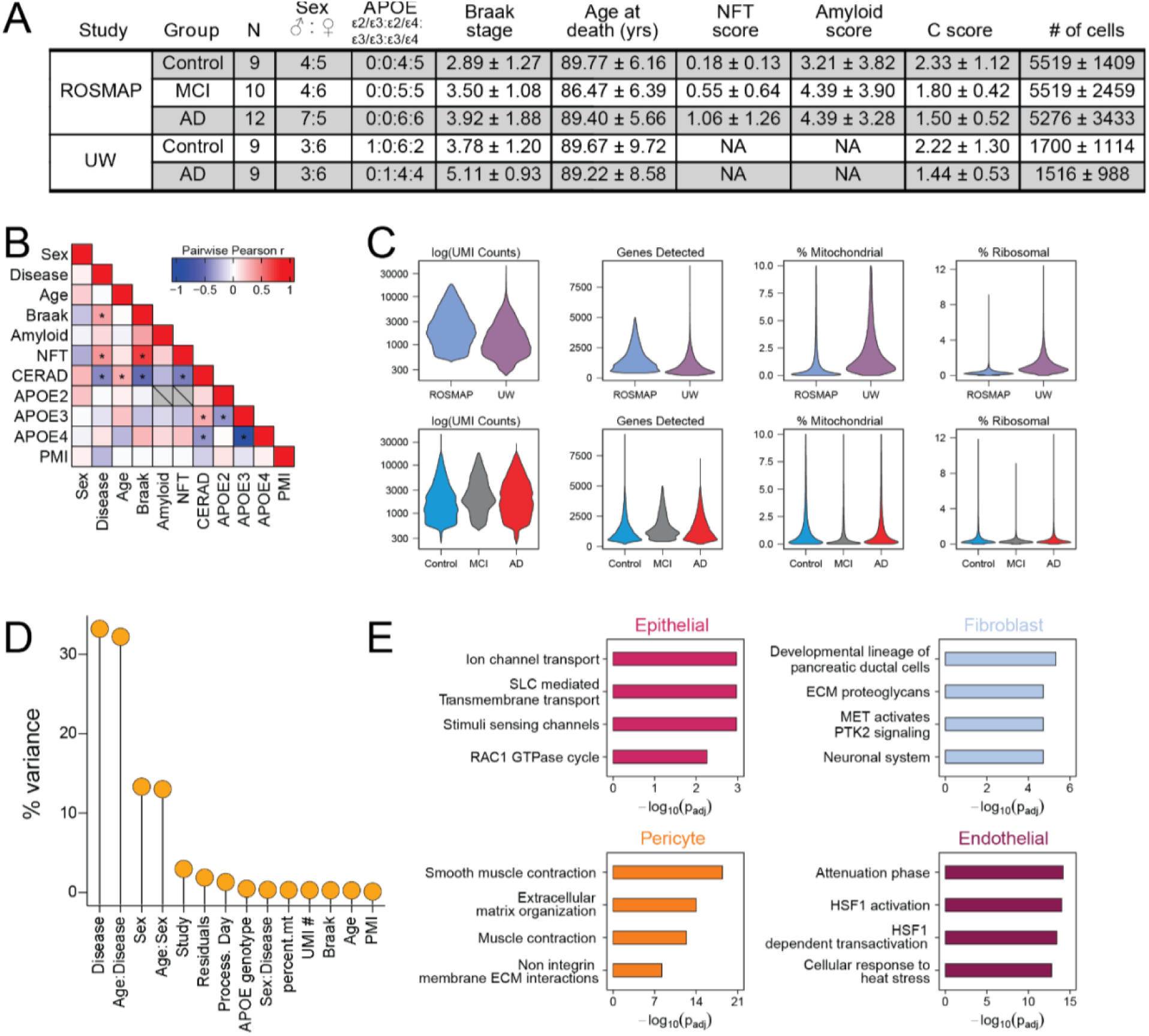
Cohort characteristics and quality control metrics for snRNA-seq, related to Figure 1. (A) Summary table of donor demographics and pathological scores by study and disease group. (B) Pairwise Pearson correlation matrix of clinical and demographic variables. * = p < 0.05. NFT, neurofibrillary tangles; APOE2/3/4, number of respective APOE alleles; PMI, post-mortem interval. (C) Violin plot of snRNA-seq quality metrics by study (top) and disease status (bottom). (D) Percent variance in gene expression explained by technical and demographic covariates. (E) Gene set enrichment analysis for cell type marker genes.

**Supplemental Figure 2:**
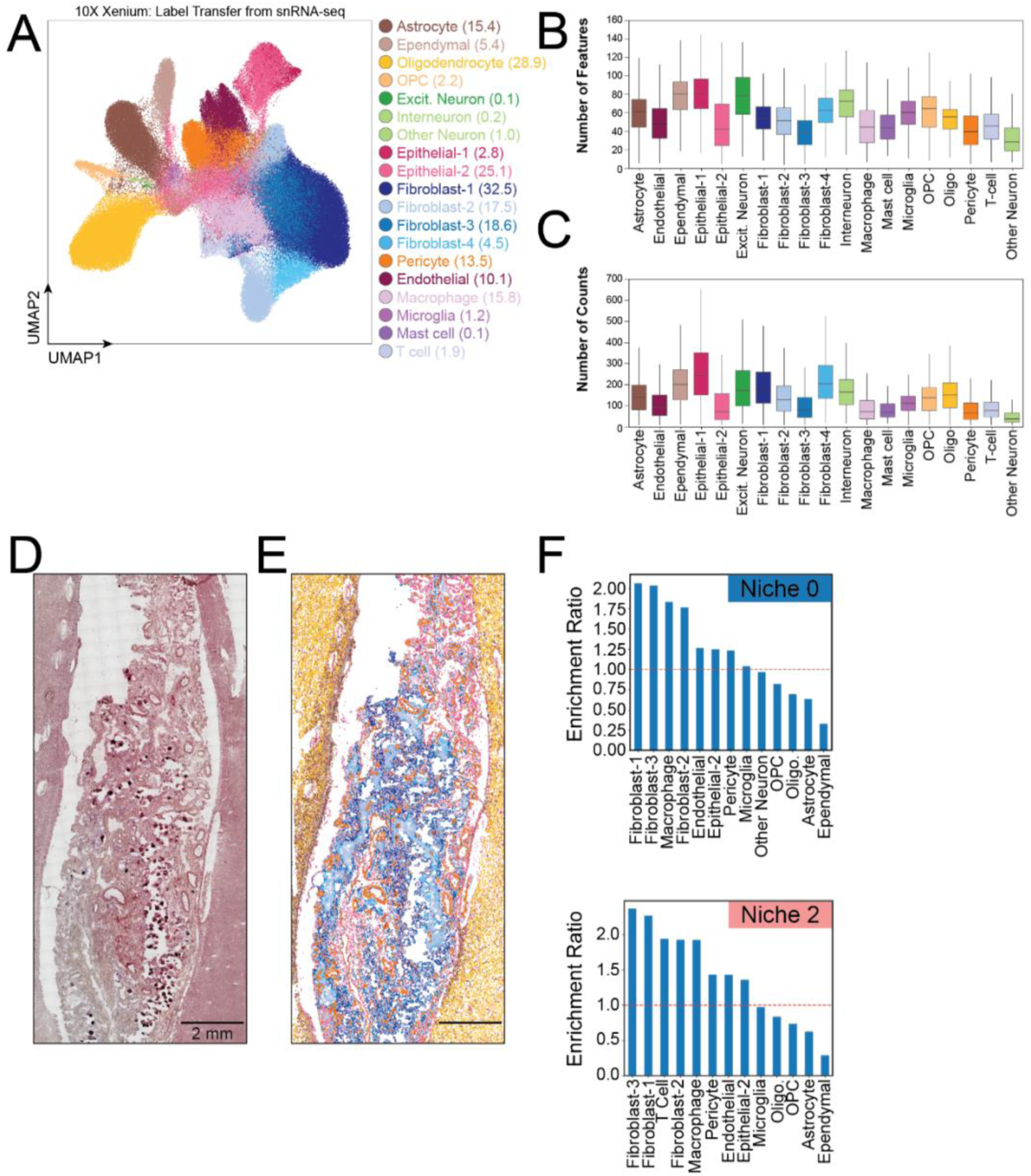
Spatial transcriptomics quality control and niche characterization, related to Figure 3. (A) UMAP visualization of Xenium data following label transfer from snRNA-seq reference. Cell type annotations and cell counts in legend. (**B-C**) Box plots of QC metrics by cell type. (**D**) H&E-stained section of human ChP tissue used for spatial transcriptomics. (**E**) Spatial map of the same tissue section colored by cell type assignment, colored by legend in (A). (**F**) Cell type enrichment ratio for spatial niches outside the ChP.

**Supplemental Figure 3.**
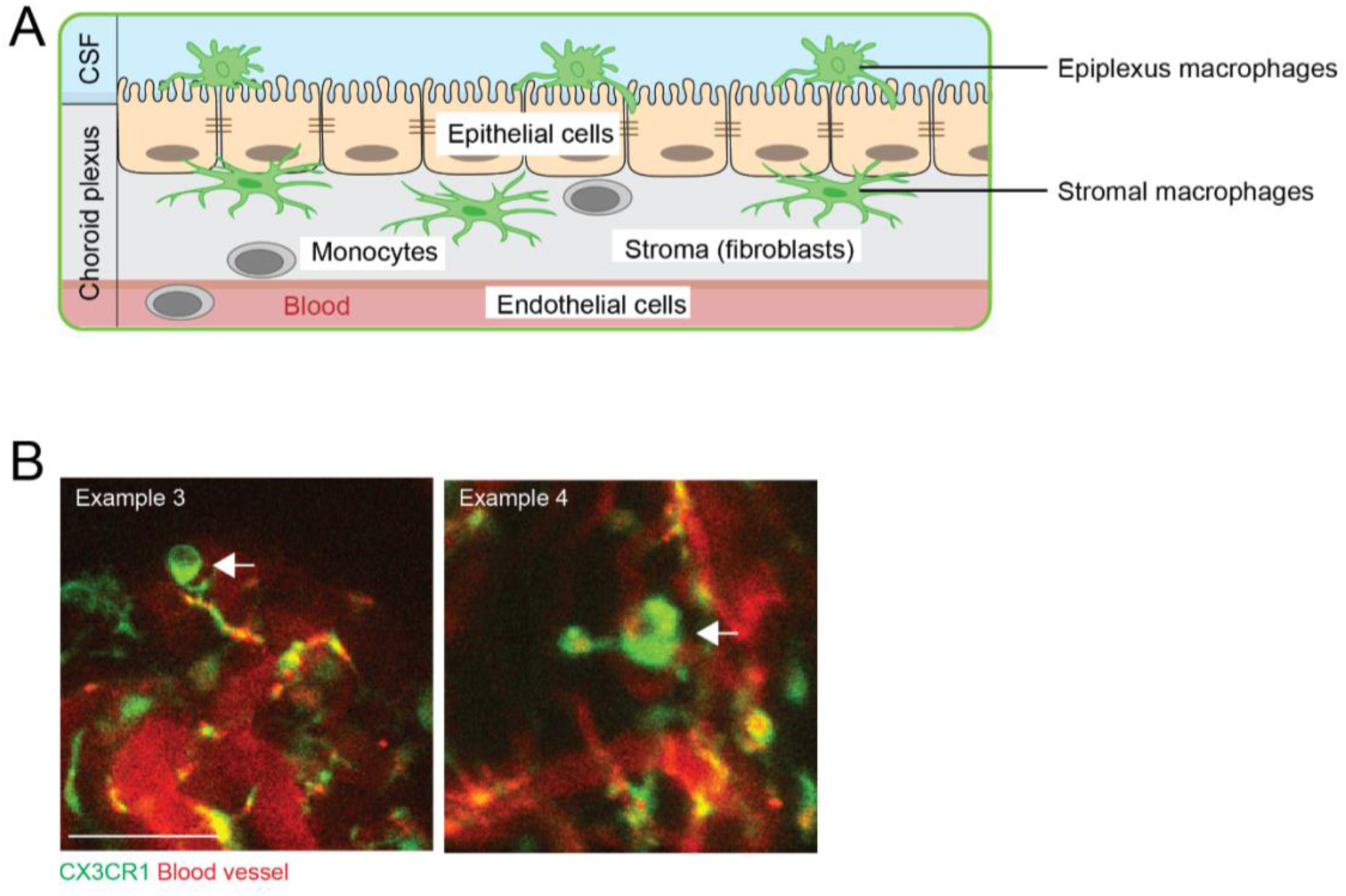
Related to Figure 4. (A) Schematics showing the anatomy of the ChP, the connection of epithelial cells by tight junctions, and the positions of epiplexus and stromal macrophages. (B) Two additional examples showing macrophages with large vacuoles, as shown in Figure 4J. Scale = 100 µm.

**Supplemental Figure 4.**
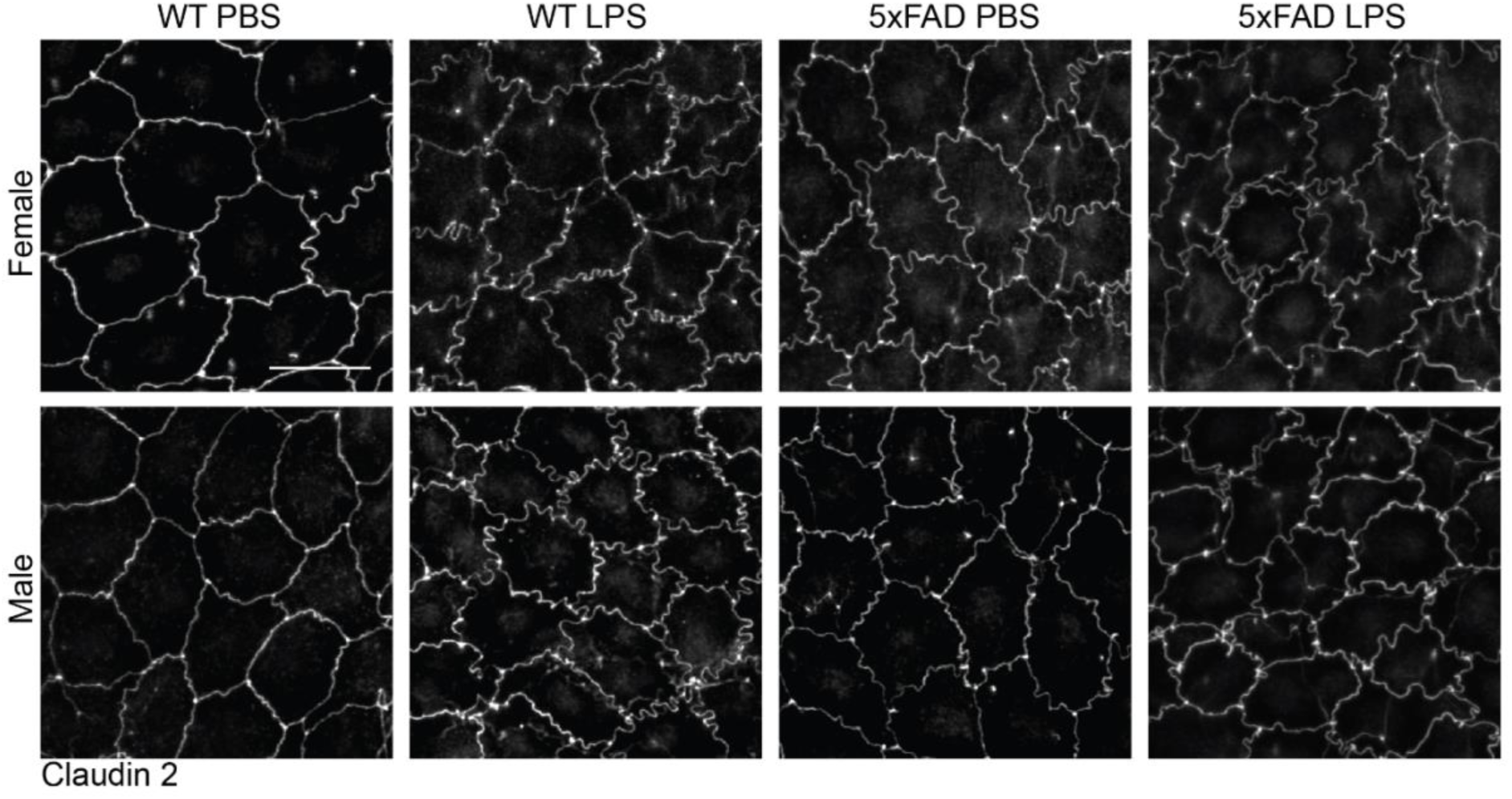
Related to Figure 5. Disruption of epithelial tight junctions in both WT and 5xFAD mice by LPS at 24 hrs. Scale = 20 µm.

**Supplemental Figure 5.**
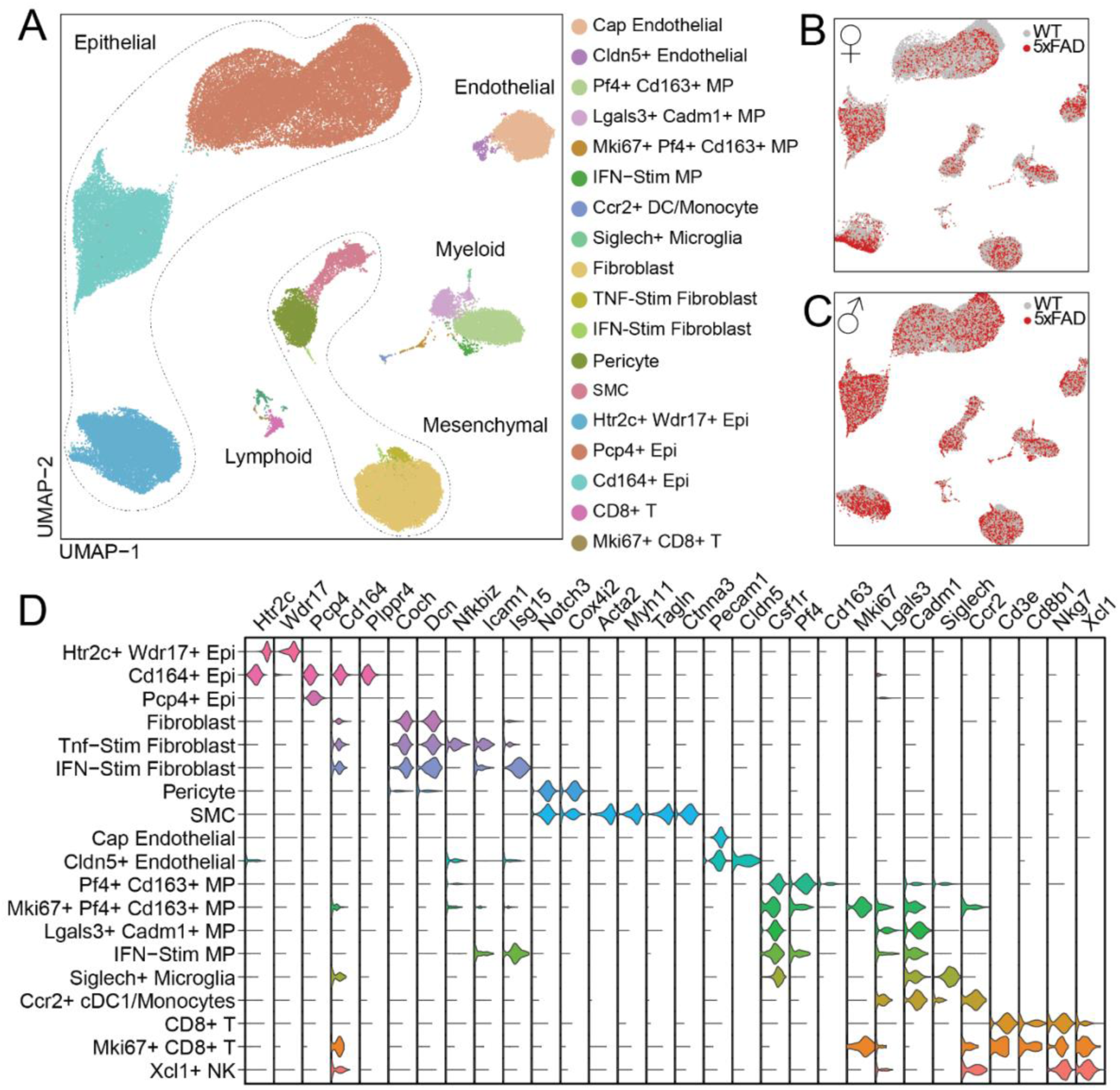
Supporting data of single-cell RNAseq of 5xFAD mice ChP at 6 months, related to Figure 6. (**A-C**) UMAP embeddings of cell states (A) and genotypes (B, females; C, males) in the 5xFAD mouse ChP at 6 months old. (**D**) Violin plot of marker gene expression across ChP-resident cell states/end-clusters.

## LIST OF SUPPLEMENTAL FILES

**Supplemental Data 1:** Patient metadata and cell type marker genes for human snRNA-seq

**Supplemental Data 2:** Patient demographic information, methodology, and statistics of histopathology analyses

**Supplemental Data 3:** Cell type marker genes for mouse ChP scRNA-seq

**Supplemental Video 1:** 5xFAD mice ChP epiplexus macrophages had abnormal morphology and some contained blood-borne fluorescent dye. Green: CX3CR1+ macrophages. Red: 70kDa Dextran-TexasRed delivered I.P. to label blood vessels. Scale = 100 µm.

**Supplemental Video 2:** 5xFAD mice ChP had epiplexus macrophages with long processes and reduced motility. Green: CX3CR1+ macrophages. Red: 70kDa Dextran-TexasRed delivered I.P. to label blood vessels. Scale = 100 µm.

**Supplemental Video 3:** 5xFAD mice ChP had macrophages forming large clusters near blood vessels. Green: CX3CR1+ macrophages. Red: 70kDa Dextran-TexasRed delivered I.P. to label blood vessels. Scale = 100 µm.

**Supplemental Video 4:** 5xFAD mice ChP had more CCR2+ cells with ramified macrophage-like morphology. Green: CCR2+ myeloid cells. Red: 70kDa Dextran-TexasRed delivered I.P. to label blood vessels. Scale = 100 µm.

## Notes

### Summary of Updates

Fixed missing citations and mislabeled Supplemental Data 3.

